# Retinoic acid signaling modulation guides *in vitro* specification of human heart field-specific progenitor pools

**DOI:** 10.1101/2022.05.30.494027

**Authors:** Dorota Zawada, Jessica Kornherr, Anna B. Meier, Gianluca Santamaria, Tatjana Dorn, Daniel Ortmann, Mark Lachmann, Mariaestela Ortiz, Stephen C. Harmer, Muriel Nobles, Andrew Tinker, Roger A. Pedersen, Phillip Grote, Karl-Ludwig Laugwitz, Alessandra Moretti, Alexander Goedel

## Abstract

Cardiogenesis relies on the precise spatiotemporal coordination of multiple progenitor populations. Understanding the specification and differentiation of these distinct progenitor pools during human embryonic development is crucial for advancing our knowledge of congenital cardiac malformations and designing new regenerative therapies. By combining genetic labelling, single-cell transcriptomics, and *ex vivo* human-mouse embryonic chimeras we uncovered that modulation of retinoic acid signaling instructs human pluripotent stem cells to form heart field-specific progenitors with distinct fate potentials. In addition to the classical first and second heart fields, we observed the appearance of juxta-cardiac field progenitors giving rise to both myocardial and epicardial cells. Applying these findings to stem-cell based disease modelling we identified specific transcriptional dysregulation in first and second heart field progenitors derived from stem cells of a patient with hypoplastic left heart syndrome. This highlights the suitability of our *in vitro* differentiation platform for studying human cardiac development and disease.

## INTRODUCTION

The heart is the first organ that forms during embryogenesis and its functionality depends on the proper spatiotemporal assembly of multiple progenitor cell populations that ultimately give rise to the diverse cell types within the specialized cardiac structures.

Lineage tracing and clonal cell analyses in mice have identified two main sources of cardiovascular progenitors in the developing mesoderm, known as the first and second heart fields (FHF and SHF), which show differential contribution to specific heart compartments^1^. A growing body of evidence suggests that the specification of FHF and SHF progenitors already occurs during gastrulation depending on their position within the primitive streak, which impacts the signaling cues they receive^2,3^. Moreover, besides having discrete molecular signatures at primitive streak stages, progenitors that later contribute to the ventricles or to the outflow tract (OFT) and atria leave the streak at different time points and form distinct regions of the cardiogenic mesoderm^4^. The heterogeneity of the heart fields is further highlighted by the recent discovery of a multipotent cardiac progenitor that generates cardiomyocytes (CMs) of the left ventricle (LV), atria and atrioventricular canal (AVC) as well as cells of the epicardium. This so-called juxta-cardiac field (JCF) resides in a distinct region within the FHF in close proximity to the extraembryonic tissue^5,6^.

The patterning of the primitive streak within the embryo occurs in response to specific signaling cues including Wnt, BMP, and Activin signaling^7^. Stimulation and inhibition of these signaling pathways in a temporally controlled manner can be used to direct *in vitro* differentiation of human pluripotent stem cells (hPSCs) into a posterior or anterior primitive streak-like fate, with cardiovascular progenitor cells arising from the latter^8^. Formation of the anterior-posterior boundaries of the cardiogenic mesoderm is, in part, controlled by retinoic acid (RA) signaling^9^, which restricts the border of the anterior SHF (aSHF – ultimately giving rise the outflow tract (OFT) and the right ventricle (RV))^10^ while maintaining its posterior component (posterior SHF (pSHF) – which generates the atria and the sinus venosus). Posteriorization of the SHF through RA signaling has been utilized during cardiac differentiation of hPSCs to promote the formation of atrial CMs and epicardial cells^12,13^. Notably, addition of low-dose RA during early mesoderm formation results in the generation of hPSC-derived cardiovascular progenitors that can self-organize into 3-dimensional (3D) structures comprising of CMs and endocardial cells, as reported recently^11^. On a transcriptional level, these progenitors resemble murine FHF cells, which are the first to migrate from the primitive streak during mid-streak stages, form the primitive heart tube, and later give rise to the LV ^4,11^. Such effect of RA during early *in vitro* cardiogenesis could reflect its inhibitory role on aSHF development providing a permissive environment for FHF lineage commitment, which has yet not been investigated in detail.

Our study aimed at systemically defining how RA impacts human cardiovascular progenitor specification during early mesoderm formation. We approached this by combining genetic labelling, transcriptomics, *ex vivo* human-mouse embryonic chimeras, and *in vitro* disease modelling. Our findings indicate that a wide spectrum of mesoderm-derived cardiovascular progenitor cells including progenitors of the JCF can be differentiated from hPSCs by modulating RA signaling in early mesoderm-like cells. These progenitors contribute to the expected cardiac structures when injected into the cardiac crescent of *ex vivo* cultured mouse embryos, highlighting the similarity to their native counterparts. Moreover, they allowed us to uncover heart field-specific differentiation defects in an *in vitro* disease model of hypoplastic left heart syndrome (HLHS), a congenital heart disease that primarily affects the LV and left ventricular outflow tract (LVOT). Overall, this study presents a versatile toolbox for the generation of defined cardiovascular progenitor pools from hPSCs and shows its applicability for studying chamber-specific heart disease.

## RESULTS

### RA signaling and cardiovascular progenitor specification

To investigate the influence of RA dosage and timing on the appearance and characteristics of early human cardiovascular progenitors in the cardiogenic mesoderm, we utilized a growth-factor based protocol for the directed differentiation of hPSCs towards CMs (Fig. 1a). In the absence of RA, mesoderm induction (*TBXT*) was detected within the first 24 hours of differentiation, followed by the expression of early cardiac progenitor markers (*NKX2.5*, *ISL1*) at day 3, shortly after Wnt signaling inhibition (Fig. 1b). Concomitant upregulation of *BMP4*, *FGF10*, *CXCR4*, and *LGR5* suggested that these cells adopted a fate resembling cardiovascular progenitors of the aSHF (Fig. 1c)^14–17^. Interestingly, addition of RA from day 1.5 to day 5.5 resulted in a dose-dependent downregulation of aSHF markers at the progenitor state (d4-5) and an upregulation of genes related to a posterior fate, such as *WNT2* and *HOXB1*, suggesting a posteriorization of the cardiac progenitor pool^15,18,19^ (Fig. 1d). This is in line with observations during cardiac development in the mouse, where RA is critical in establishing the anterior-posterior axis within the heart fields^20^. Notably, we also detected a strong upregulation of *TBX5*, which at the equivalent stage *in vivo* marks cardiac progenitors of the murine FHF^15^, as well as *THBS4*, which is highly enriched in FHF progenitor cells in the mouse embryo at embryonic day 7.75 (E7.75)^6^ (Fig. 1d). During further differentiation, we observed earlier and higher expression of sarcomeric genes such as *TNNT2* and *MYL3* in presence of RA compared to differentiation without RA, which was accompanied by a faster downregulation of *ISL1* (Fig. 1e, Supplementary Fig. 1a). Concordantly, beating foci emerged on average two days earlier (Fig. 1f). Notably, rapid downregulation of *ISL1* expression and rapid generation of functional CMs are among the characteristic features of FHF cardiovascular progenitors, which are necessary to form the primitive heart tube *in vivo*^4^.

**Figure 1:**
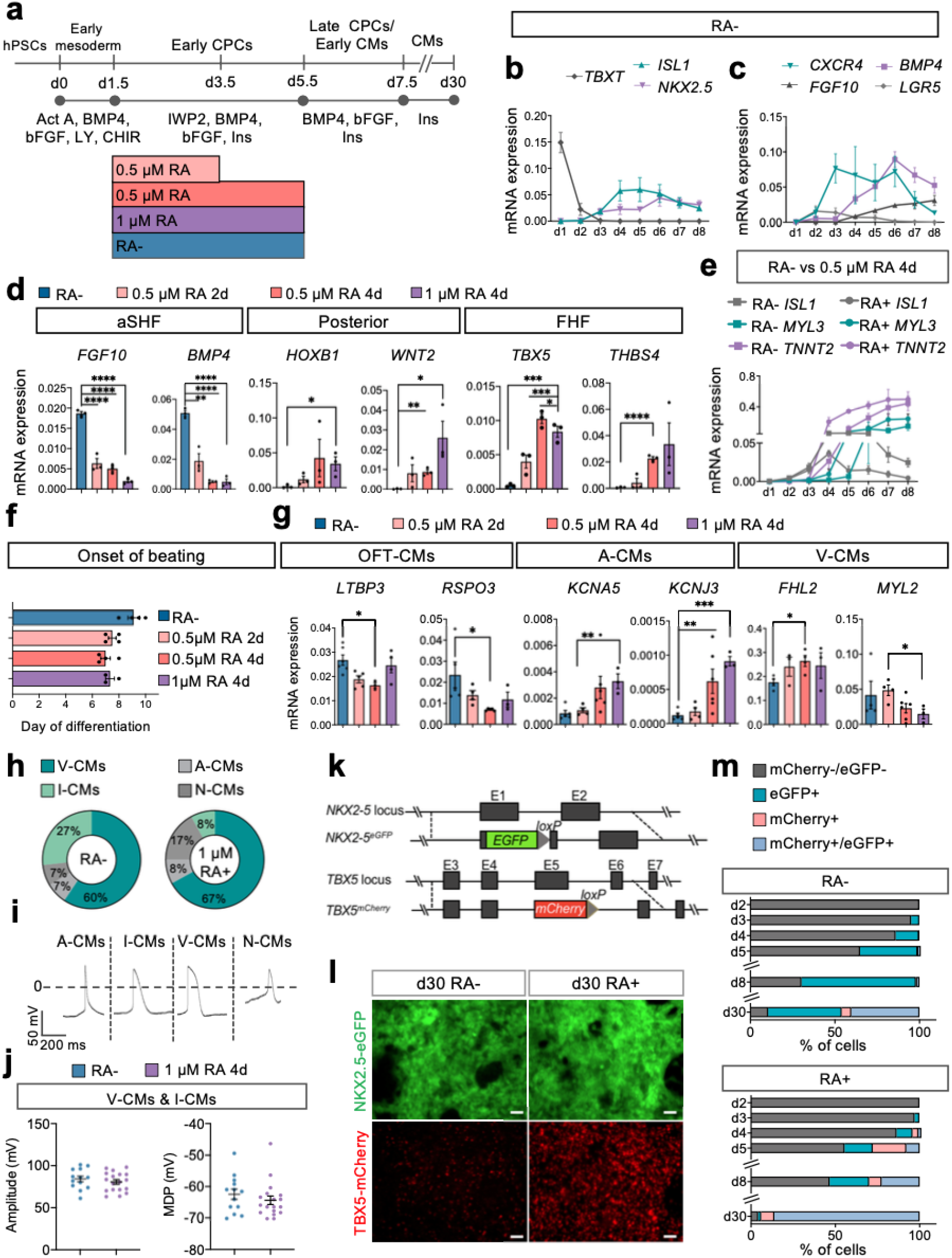
RA signaling modulation impacts cardiovascular progenitor specification. **(a)** Schematic representation of the protocol used to differentiate human pluripotent stem cells (hPSCs) into cardiomyocytes (CMs) through defined steps of early mesoderm and cardiovascular progenitor cells (CPCs) without retinoic acid (RA-) or with RA added at the indicated dosages and times. Act A: Activin A; CHIR: CHIR-99021; LY: LY-29004; Ins: insulin. **(b,c)** Time course of mRNA expression of early mesoderm and CPC markers **(b)** and anterior second heart field (aSHF) markers **(c)** during differentiation without RA. Data are mean ± SEM. n = 3 differentiations/time point. mRNA expression relative to *GAPDH*; *p<0.05, **p<0.005, ***p<0.001 (unpaired two-tailed *t*-test), unless otherwise indicated. **(d)** mRNA expression of key anterior second heart field (aSHF) markers, posteriorization markers, and first heart field (FHF) markers at the CPC stage in the indicated differentiation conditions. Samples analysed at d4 (for *HOXB1*) or d5 (for other genes). Data are mean ± SEM; n = 3 differentiations/time point. **(e)** Time course of mRNA expression of CPCs and early CM markers during RA- and RA+ (0.5 μM 4d) differentiation. Data are mean ± SEM. n = 3 differentiations/time point. **(f)** Day of the onset of spontaneous beating in the indicated differentiation conditions. Data are mean ± SEM; n = 3-4 differentiations. **(g)** mRNA expression of markers of outflow tract CMs (OFT-CMs), atrial CMs (A-CMs) and ventricular CMs (V-CMs) relative to *TNNT2* and *GAPDH* at d30 of the indicated differentiation conditions. Data are mean ± SEM; n ≥ 3 differentiations/time point. **(h)** Percentage of cardiomyocyte subtypes at d30 of differentiation with 1 μM RA for 4d or without RA (RA-) as defined by the ratio of action potential duration at 50% and 90% repolarization determined by whole-cell voltage clamp recording. I-CMs: Intermediate ventricular cardiomyocytes; V-CMs: Ventricular cardiomyocytes; A-CMs: atrial cardiomyocytes; N-CMs: nodal cardiomyocytes. **(i)** Representative action potential traces of indicated cardiomyocyte subtypes at day 30. **(j)** Amplitude (left) and Maximum Diastolic Potential (MDP, right) of action potentials in cardiomyocytes at d30 of differentiation with 1 μM RA for 4d or without RA (RA-). Data are mean ± SEM. For 1 μM RA for 4d N = 18 cells; for RA-N = 13 cells from 2 differentiations. **(k)** Schematic representation of the genetic modifications in the TBX5^mCherry^ and NKX2.5^eGFP^ hESC double reporter cell line (ES03 TN). **(l)** Representative live images of cells expressing eGFP (NKX2.5) or mCherry (TBX5) at d30 of differentiation with 0.5 μM RA for 4d (RA+) or without RA (RA-). Scale bar = 50 μm. **(m)** Percentage of cells expressing mCherry (TBX5) and eGFP (NKX2.5) at the indicated days of differentiation with 0.5 μM RA for 4d (RA+) or without RA (RA-) as determined by live flow cytometry. Data are mean; n ≥ 3 differentiations.

At day 30, expression levels of *TNNT2* as well as the percentage of cTnT+ cells were comparable between differentiations with and without RA, suggesting efficient CM differentiation in both conditions (Supplementary Fig. 1b-c). Without RA, cells expressed high levels of ventricular CM genes such as *MYL2*, *IRX4*, and *IRX3*, while atrial CM markers such as *KCNJ3* and *KCNA5* were absent (Fig. 1g; Supplementary Fig. 1c). In addition, these cells expressed transcripts typical of CMs found in the OFT, such as *LTBP3* and *RSPO3*^16^ (Fig. 1g), providing further evidence that this differentiation condition induces an aSHF-like fate. Upon RA addition, we detected a dose-dependent increase in expression of *FHL2* (Fig. 1g), which was reported to be enriched in left ventricular CMs of adult human hearts^21^. Genes associated with CMs of the OFT were downregulated upon RA treatment across the dosage range, while atrial CM markers marginally increased with higher RA exposure (Fig. 1g). In line with our findings on the transcriptomic level, electrophysiological examination showed that the majority of CMs obtained with and without RA were of ventricular rather than atrial or nodal identity (Fig. 1h,i; Supplementary Table 1). Without RA, over a quarter of cells showed “intermediate” action potential characteristics, suggesting that they have not fully differentiated yet (Fig. 1h,i, Table S1). However, action potential properties such as the mean diastolic potential and amplitude of the cells classified as ventricular were comparable between the two groups (Fig. 1j). Notably, even at higher doses of RA (1 μM, 4 days) the proportion of atrial cells was less than 10% (Fig. 1h), suggesting that in these differentiation conditions RA exposure does not result in atrial fate^12,22^. Instead, we did observe a slight increase in cells with a nodal-like action potential (Fig. 1h), probably reflecting a permissive environment for sinus nodal commitment^23^.

Based on these findings, we hypothesized that a time- and dose-limited exposure to RA during early mesoderm formation is not efficient in inducing a pSHF-like fate resulting in atrial CMs, but rather promotes the specification of progenitors resembling a FHF-like fate, resulting in CMs with a (left) ventricular identity. Since the intermediate dose of RA (0.5 μm, 4 days) was sufficient to suppress aSHF genes and induce FHF commitment (Fig. 1d) we decided to use this dose in further experiments.

To track the appearance of distinct cardiovascular progenitor pools and enable enrichment of specific subpopulations for in-depth analysis, we generated a double reporter human embryonic stem cell (hESC) line expressing eGFP and mCherry under the control of the endogenous *NKX2.5* and *TBX5* locus, respectively (ES03 TN cell line, Fig. 1k). We confirmed proper reporting of the fluorescent markers using live imaging (Fig. 1l), RT-qPCR, and immunofluorescence staining (Supplementary Fig. 1d,e). During differentiation, the modified hESC line displayed a slight reduction in expression levels of *NKX2.5* and *TBX5* compared to the parental line (Supplementary Fig. 2a). However, the expression patterns of key cardiac differentiation genes and the percentage of cTnT+ cells were comparable between the reporter and parental line as well as an unrelated human induced pluripotent stem cell (hiPSC) line (Supplementary Fig. 2a-c). Without addition of RA, eGFP^+^ cells (NKX2.5^+^) appeared from day 3 while mCherry^+^ (TBX5^+^) cells were virtually absent until day 8 (Fig. 1m; Supplementary Fig. 2d’). eGFP^+^ cells maintained ISL1 expression while progressively upregulating the CM marker cTnT from day 5 (Supplementary Fig. 2e). With addition of RA, eGFP^+^ cells were first detected at day 3 and both mCherry^+^ and eGFP^+^/mCherry^+^ populations emerged from day 4 (Fig. 1m; Supplementary Fig. 2d’’). In this condition, we observed a rapid downregulation of ISL1 and upregulation of cTnT in all populations (Supplementary Fig. 2e), again suggestive of a FHF-like fate.

### RA signaling patterns the cardiogenic mesoderm

To gain further insights into the formation of cardiac progenitors in response to limited RA exposure, we performed single-cell RNA sequencing (scRNA-seq) on early mesodermal cells at day 1.5 (right before the addition of RA) as well as on individual cardiovascular progenitor populations at day 4.5 (Fig. 2a). Using flow cytometry-based sorting, we isolated eGFP^+^ (NKX2.5) cells obtained in the absence of RA and the three populations with differential eGFP (NKX2.5) and mCherry (TBX5) expression emerging in the presence of RA (Fig. 2a). Low dimensionality embedding (UMAP) of all cells merged resulted in the formation of five distinct groups: early mesodermal cells (day 1.5), endothelial/endocardial progenitor cells (E-PCs), endodermal cells, and cardiovascular progenitor cells derived in the absence (CPC-RA-) or presence of RA (CPC-RA+) (Fig. 2b; Supplementary Table 2).

**Figure 2:**
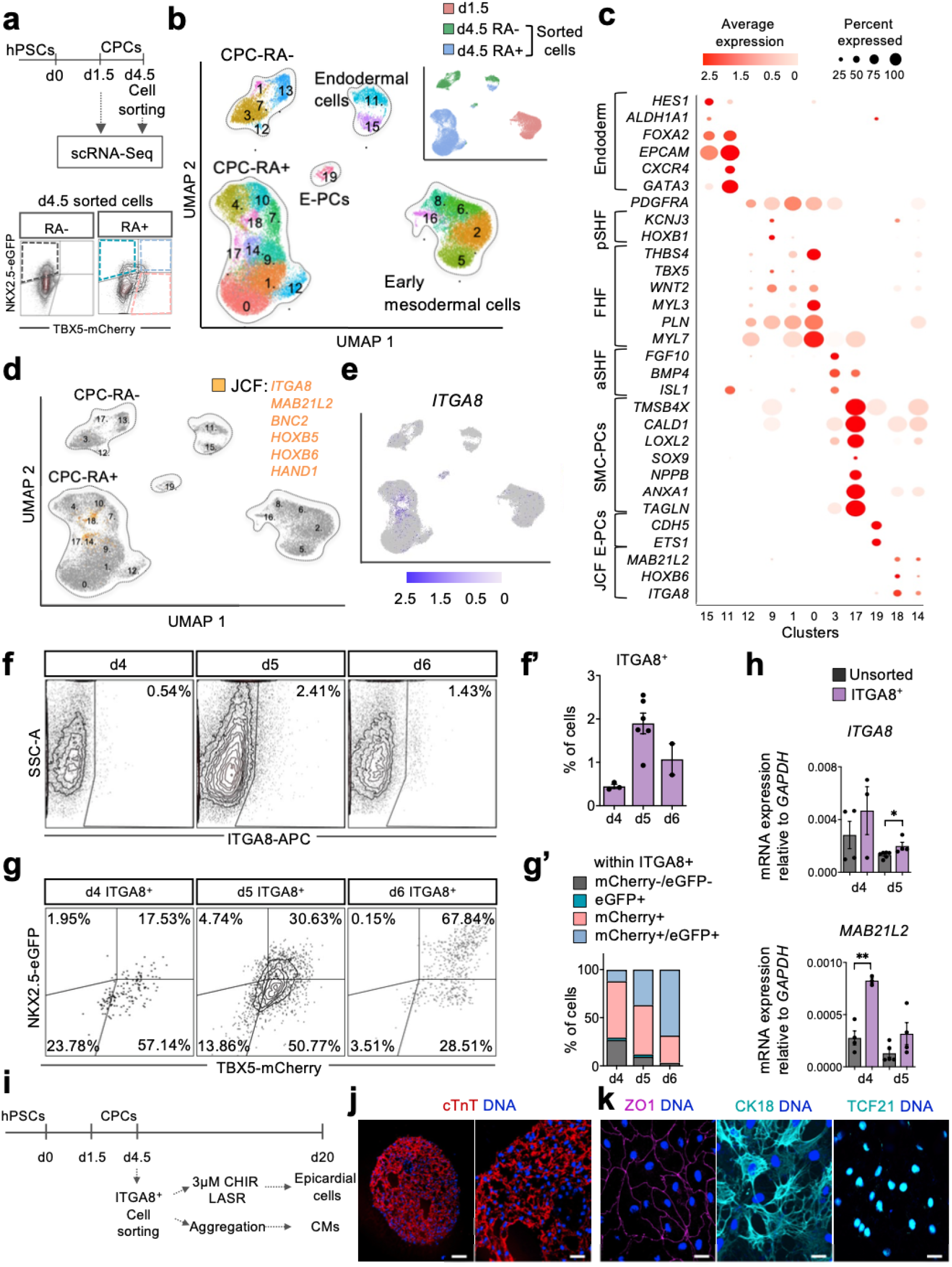
RA signaling patterns the cardiogenic mesoderm. **(a)** Top: Graphic representation of the experimental design applied for scRNA-Seq analysis at days 1.5 and 4.5. Bottom: Representative flow cytometry plots showing the gating strategy for sorting cells at d4.5. Cells differentiated without RA (RA-) were isolated based on expression of eGFP (NKX2.5); cells differentiated with 0.5 μM RA for 4d (RA+) were isolated based on expression of eGFP (NKX2.5), mCherry (TBX5), or both. **(b)** UMAP clustering of single cells captured at d1.5 and d4.5; main cell types are annotated. Inset: UMAP plot showing the contribution of the indicated samples. CPCs: cardiovascular progenitor cells; E-PCs: endothelial/endocardial progenitor cells. **(c)** Dot plot showing the expression level of selected differentially expressed genes for clusters identified as 15, 11 - endoderm; 12 – early CPCs; 9 – posterior second heart field (pSHF); 1, 0 – first heart field (FHF); 3 – anterior second heart field (aSHF); 17 – smooth muscle cell progenitor cells (SMC-PCs); 19–E-PCs; 18, 14 – juxta-cardiac field (JCF). **(d)** Feature plot showing cells co-expressing key JCF markers in orange. **(e)** Feature plot showing the expression of *ITGA8*. **(f)** Representative plots and **(f’)** quantification of live flow cytometry analysis of cells expressing ITGA8 (APC) during differentiation with 0.5 μM RA for 4d. Data are mean ± SEM; n = 2-5 differentiations. **(g)** Representative plots and **(g’)** quantification of live flow cytometry analysis of cells expressing mCherry (TBX5) and eGFP (NKX2.5) within the ITGA8+ (APC) population during differentiation with 0.5 μM RA for 4d. Data are mean; n = 2-5 differentiations. **(h)** mRNA expression of *ITGA8* and *MAB21L2* relative to *GAPDH* in cells FACS sorted for ITGA8 (APC) and unsorted cells during differentiation with 0.5 μM RA for 4d. Data are mean ± SEM; n ≥ 3 differentiations/time point; *p<0.05, **p<0.005, ***p<0.001 (unpaired two-tailed *t*-test). **(i)** Schematic representation of the experimental timeline for FACS sorting of ITGA8^+^ cells followed by epicardial and myocardial differentiation. **(j,k)** Representative images of cells expressing myocardial **(j)** and epicardial markers **(k)** 15 days after replating/reaggregation of ITGA8^+^ sorted progenitors at d4.5 of differentiation with 0.5 μM for 4d. For (j) scale bar = 100 μm (left) and 30 μm (right). For (k) scale bar = 20 μm.

Cells at day 1.5 represented a homogenous cell population expressing mesodermal genes such as *TBXT* and *MESP1* (Supplementary Fig. 3a). Integration of this data with a recently published scRNA-seq dataset of a gastrulating human embryo^24^ showed that these cells correspond to an early mesoderm population that is in the process of emerging from the primitive streak (emergent mesoderm, Supplementary Fig. 3b). The expression of genes such as *EOMES*, *GSC*, and *MIXL1* suggested that these cells match a mid/anterior primitive streak fate^8^ (Supplementary Fig. 3c). Notably, we observed high expression of the cytochrome member *CYP26A1*, which has been reported to mark a subpopulation of the cardiogenic mesoderm prone to giving rise to ventricular cardiomyocytes^12^ (Supplementary Fig. 3d).

The cardiovascular progenitors derived in the absence of RA (CPC-RA-) separated into two main clusters based on their proliferative state (cluster 3, non-proliferative; cluster 13, proliferative; Fig. 2b; Supplementary Fig. 4a), both showing expression of aSHF genes such as *ISL1*, *FGF10*, and *BMP4* (Fig. 2c; Supplementary Fig. 4b). Two small additional clusters were shared with the RA+ condition: cluster 12 was characterized by high expression of *PDGFRA*, indicating undifferentiated cardiovascular progenitor cells, and cluster 17 was enriched in genes related to smooth muscle cells (SMCs) such as *TAGLN*, *NPPB*, *LOXL2*, *TMSB4X*, and *CALD1* (Fig. 2b,c; Supplementary Fig. 4c).

In the presence of RA, the cardiovascular progenitor cells (CPC-RA+) formed several clusters (Fig. 2b). Clusters 0 and 1 showed expression of *THBS4* as well as cardiac sarcomere protein genes such as *MYL3*, *MYL7*, and *PLN*, most likely resembling progenitors of the FHF that are in the process of converting to CM (Fig. 2c). Cluster 9 was characterized by expression of posterior transcripts such as *WNT2* and *HOXB1*^15^, possibly representing a small fraction of pSHF cells (Fig. 2c). This cluster was mainly derived from TBX5^+^/NKX2.5^−^ progenitors (~75%; Supplementary Fig. 4c). Clusters 4, 7, and 10 represented the proliferative equivalents of clusters 0, 1, and 9, respectively (Supplementary Fig. 4a).

Notably, among the CPC-RA+ we observed two small additional clusters (cluster 14 and its proliferative counterpart cluster 18) that showed a gene expression profile (*HAND1*, *MAB21L2*, *HOXB6*, *HOXB5*, *BNC2*) resembling that of the recently described JCF, which is a distinct subset of the FHF in mice (Fig. 2c,d). Progenitors residing in this field have the potential to contribute to the cardiomyocytic as well as the epicardial lineage^6^. This suggests that a similar population could exist during human cardiogenesis.

### ITGA8 allows isolation of human progenitors corresponding to juxta-cardiac field cells

To further explore this putative JCF cell population, we aimed at identifying a surface marker allowing the specific isolation of these cells. Comparative differential gene expression analysis between the mouse dataset of Tyser et al. (2021) and ours revealed *TNC*, *AHNAK*, and *ITGA8* as promising candidates showing co-expression with other key JCF markers (*MAB21L2*, *HOXB5*, *HOXB6*, *HAND1*, *BNC2*; Supplementary Table 3)^6^. Among these, *ITGA8* showed the most restricted expression pattern (Fig. 2e; Supplementary Fig. 4d,e). Flow cytometry analysis confirmed the presence of a small population of ITGA8^+^ cells from d4 to d6 (Fig. 2f’-f”). This population initially expressed only TBX5 (mCherry) but quickly upregulated NKX2.5 (eGFP) (Fig. 2g’-g”), which is in line with observations in the murine JCF^6^. Cells sorted based on the presence of ITGA8 showed significantly higher expression of JCF-specific markers compared to unsorted cells (*MAB21L2*, *BNC2*, *HAND1*; Fig. 2h; Supplementary Fig. 4f). After cultivating ITGA8^+^ sorted cells in myocardial differentiation conditions, we obtained cTnT+ CMs (Fig. 2i,j). On the other hand, when exposed to epicardial differentiation conditions, the cells formed a tight epithelial layer staining positive for the epithelial markers ZO1 and CK18 as well as the epicardial marker TCF21 (Fig. 2i,k). This data suggests that a bipotent JCF progenitor pool giving rise to CMs and epicardial cells may also exist during human cardiogenesis.

### CPC-RA+ and CPC-RA- give rise to distinct cardiac cell populations

Next, we aimed to characterize the descendants of the cardiovascular progenitors emerging in our system. For this, cells differentiated with and without RA were sorted at day 4.5 based on eGFP (NKX2.5) and mCherry (TBX5) expression and further cultured as 3D aggregates to allow more physiological conditions for differentiation. Aggregates were then collected at day 30 for scRNA-seq (Fig. 3a). Cells from day 30 merged with the cells from day 4.5 and 1.5 in UMAP formed three distinct groups: endoderm-derived cells, non-myocytic cardiac cells, and CMs (Fig. 3b,c; Supplementary Fig. 5a; Supplementary Table 4).

**Figure 3:**
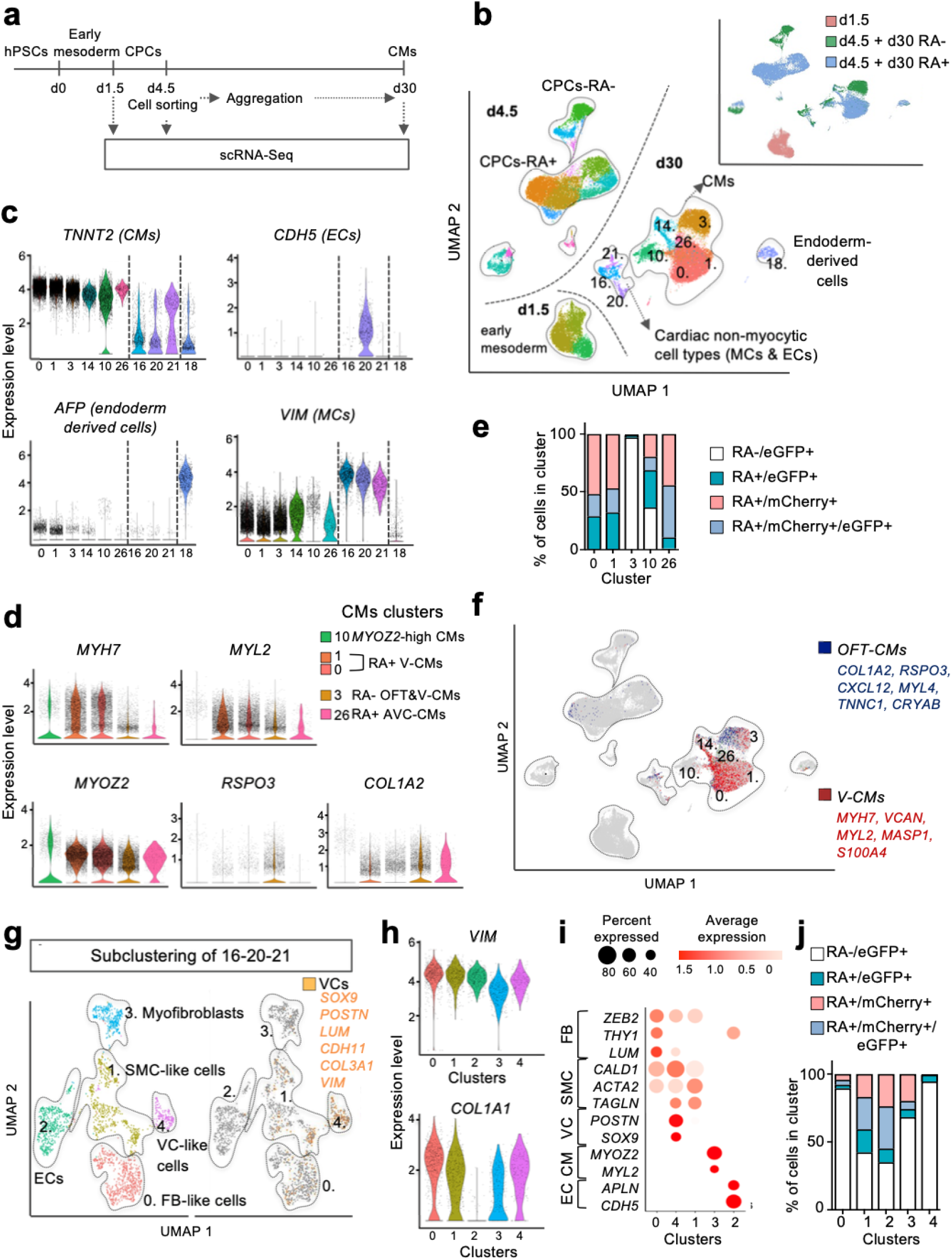
CPC-RA- and CPC-RA+ give rise to distinct cardiac cell types. **(a)** Graphic representation of the experimental design for scRNA-Seq analysis at d1.5, d4.5, and d30. At d4.5, cells differentiated without RA (RA-) were sorted based on expression of eGFP (NKX2.5); cells differentiated with 0.5 μM RA for 4d (RA+) were sorted based on expression of eGFP (NKX2.5), mCherry (TBX5), or both. Sorted cells were reaggregated in 3D and analyzed at d30. **(b)** UMAP clustering of single cells captured at d1.5, d4.5, and d30; main cell types are annotated. Inset: UMAP plot showing the contribution of the indicated samples. CMs: cardiomyocytes; MCs: mesenchymal cells; ECs: endothelial cells. **(c)** Violin plots showing the expression level of selected key cell types markers in selected clusters from (b). **(d)** Violin plots showing the expression level of selected key differentially expressed cardiomyocyte subtypes markers in CM clusters from (b). **(e)** Contribution of d30 cells derived from each sorted CPC population to the indicated CM clusters relative to all cells in the cluster: 1, 0 – ventricular cardiomyocytes (V-CMs); 3 – outflow tract and ventricular cardiomyocytes (OFT & V-CMs); 10 – *MYOZ2*-high CMs; 26 – atrioventricular canal cardiomyocytes (AVC-CMs). **(f)** Feature plot showing in red cells co-expressing top differentially expressed genes defining V-CMs clusters of Cui et al. (2019) and confirmed with specific expression in V-CMs clusters of Asp et al. (2019: online search tool); in blue cells co-expressing genes defining OFT-CMs cluster of Li et al (2016). **(g)** Left: UMAP subclustering of cardiac non-myocytic clusters (16, 20, 21) at d30; right: feature plot showing in yellow cells co-expressing key markers of valvular cells (VC). **(h)** Violin plots showing the expression level of *VIM* and *COL1A1* in clusters from (g). **(i)** Dot plot showing the expression level of selected differentially expressed genes for the subclusters shown in (g). **(j)** Contribution of d30 cells derived from each sorted cell population to subclusters shown in (g) relative to all cells in the cluster.

The CMs separated into five main groups: ventricular-like CMs, OFT-like CMs, AVC-like CMs, *MYOZ2-high* CMs, and proliferating CMs (Fig. 3d; Supplementary Fig. 5a). The *MYOZ2*-high CMs, which were present in both differentiation conditions (CPC-RA- and CPC-RA+; Fig. 3d,e), most likely represent the *MYOZ2*-enriched CM subset that has been described in the LV and RV of human and mouse hearts^25,26^. The identity of the OFT-like and ventricular-like CMs was confirmed on a transcriptome-wide level using enrichment analysis for gene signature scores derived from human and mouse *in vivo* heart samples^27,28^ (Fig. 3f). OFT-like CMs were almost exclusively found as derivatives of CPC-RA-. Within cluster 3, they were located to the left-hand side of the cluster while the CPC-RA- ventricular-like CMs were located on the right. OFT-like CMs expressed markers of human conoventricular CMs, such as *BMP2* and *RSPO3*^16^, as well as SMC markers (e.g., *ACTA2*, *TAGLN*, *COL1A2*) and only low expression levels of *MYL2*, which is in line with the transcriptional profile of OFT-CMs in mice^29^ (Fig. 3d; Supplementary Figs. 5b and 6). Moreover, they did not express *TBX5*, which is consistent with data from human fetal tissue showing absence of TBX5 in OFT structures^30^ (Supplementary Fig. 6).

Subclustering of the non-myocytic cells, which had in common high expression levels of *VIM* (Fig. 3c, 3h), revealed the presence of endothelial/endocardial-like cells (*APLN*, *CDH5*), smooth muscle-like cells (*TAGLN*, *ACTA2*, *CALD1*), fibroblast-like cells (*COL1A1*, *THY1*, *LUM*), myofibroblasts (*MYOZ2*, *MYL2*), and valvular-like cells^31–33^ (*SOX9*, *POSTN*; Fig. 3g-i; Supplementary Table 5). In addition, we detected cells expressing genes associated with endothelial to mesenchymal transition (EndoMT), such as *ZEB2* and *LUM* (Fig. 3i). While endothelial/endocardial-like cells and smooth muscle-like cells were found as derivatives of all populations, valvular-like cells were almost exclusively derived from aSHF-like progenitors (CPC-RA-; Fig. 3j). This is in line with previous reports in the mouse showing that valve formation mainly relies on contribution from the aSHF^34,35^.

Overall, these findings confirmed that the differences observed at the cardiovascular progenitor state translated into the formation of distinct progeny reflecting FHF- vs aSHF-like fate potential.

### Fate decision trees of human cardiac progenitors

To reconstruct fate decisions trees for the various progenitor populations, we combined all time points (day 1.5, day 4.5 and day 30) for each condition (RA- and RA+). In the absence of RA, URD analysis inferred the presence of a common progenitor state between non-myocytic and myocytic fates at day 4.5 (Fig. 4a), which is in line with the multilineage potential of the cardiovascular progenitors of the aSHF^3,34^. These cells showed high expression of *ISL1* as well as *BMP4* (Fig. 4b)^36,37^. The subsequent myocytic fate acquisition (SMCs and CMs) was correlated with upregulation of *BMPER* (Fig. 4c; Supplementary Table 7), which has been shown in the mouse to impact cardiomyocyte size as well as vessel density^38^, and *NRG2* (Fig 4c; Supplementary Fig. 7a), which has been reported to regulate cardiac subtype specification^39^. In the cardiomyocytic lineage trajectory we observed early divergence of *MYOZ2*-high CMs (branch 7) from other CM-subtypes such as AVC- (*RSPO3*, *BMP2*, *TBX5*; branch 2), OFT- (*RSPO3*, *BMP2*, branch 3) or ventricular-like CMs (*MYH7*, *MLY2*; branches 1, and 4-6; Supplementary Fig. 7a). The non-myocytic fate was correlated with upregulation of *ETV2* (branch 24; Supplementary Fig. 7a), a transcription factor with a well-established role in endothelial fate decision^40^ and *NOTCH3*, a marker of fibroblast development^41^ (branch 24, and subsequent branches 23 and 11; Fig. 4c; Supplementary Table 7). Notably, we also identified branches (24, and subsequent branches 23, and 8) that were correlated with the upregulation of epidermal growth factor receptor (*EGFR*; Fig. 4c; Supplementary Fig. 7a), which is critically involved in the development of semilunar valves from aSHF-derived progenitors in the OFT in mice^42,43^.

**Figure 4:**
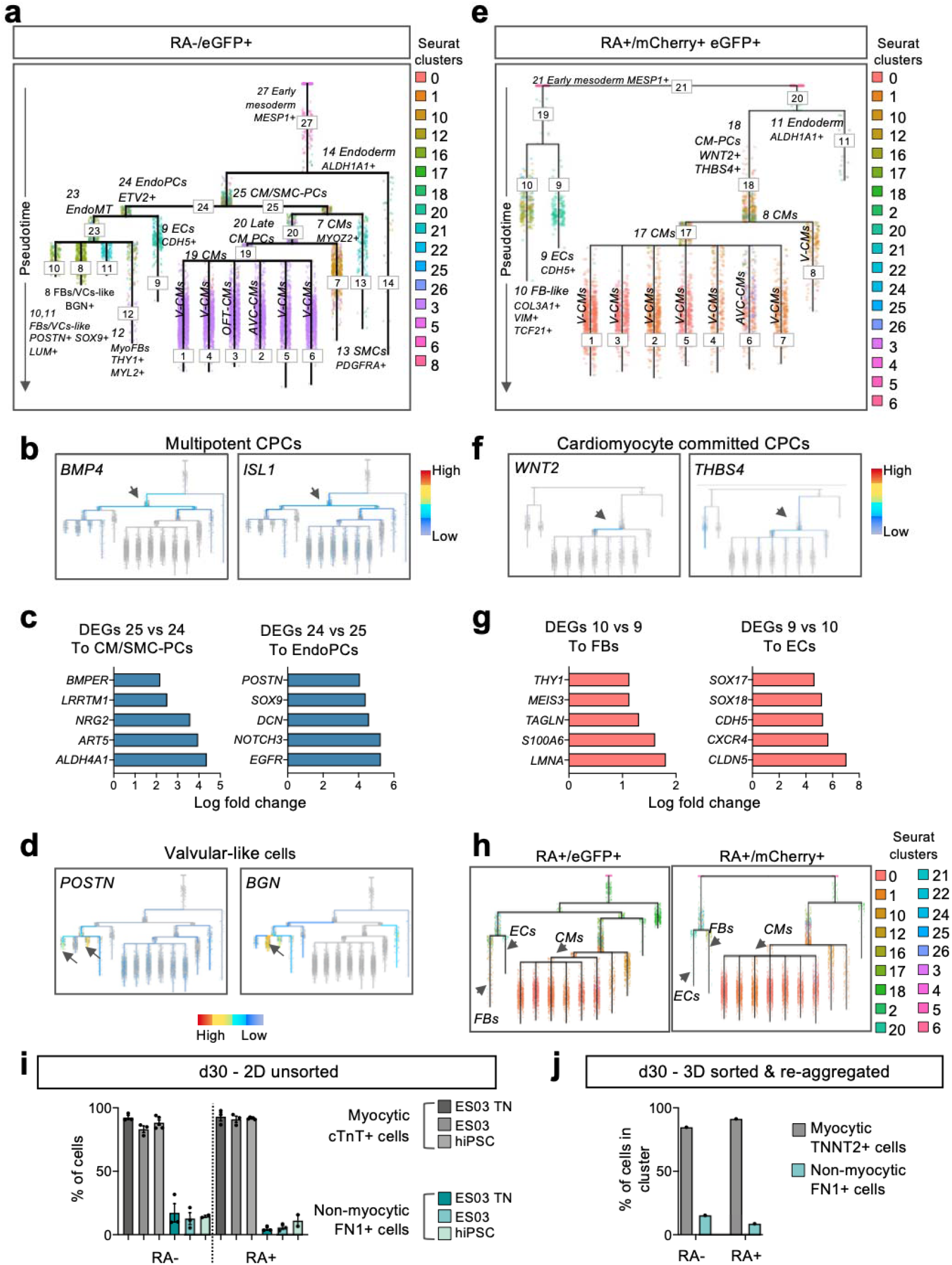
Fate decisions of human cardiac progenitors. **(a)** Dendogram showing URD inferred lineage tree of single cells at d1.5, sorted cells expressing eGFP (NKX2.5) at d4.5, and d30 cells obtained after reaggregation of cells sorted at d4.5; all from differentiation without retinoic acid (RA-). Cells from each Seurat cluster are indicated by colour. **(b)** Plots showing expression levels of multipotent CPC marker genes *BMP4* and *ISL1* on the URD branches from (a). Arrows indicate branches with relatively the highest expression of the respective genes. **(c)** Upregulated genes found differentially expressed between the indicated branches. Left: genes upregulated in branch 25 compared to 24; right: genes upregulated in branch 24 compared to 25. **(d)** Plots showing the expression levels of the valvular cell markers *POSTN* and *BGN* on the URD branches from (a). Arrows indicate branches with relatively the highest expression of the respective genes. **(e)** Dendogram showing URD inferred lineage tree of single cells captured at d1.5, sorted cells expressing both eGFP (NKX2.5) and mCherry (TBX5) at d4.5, and d30 cells obtained after reaggregation of cells sorted on d4.5; differentiation with 0.5 μM RA for 4d (RA+). Cells from each Seurat cluster are indicated by colour. **(f)** Plots showing the expression levels of cardiomyocyte committed CPC marker genes on the URD branches from (b). Arrows indicate branches with relatively the highest expression of the respective genes. **(g)** Upregulated genes found differentially expressed between the indicated branches. Left: genes upregulated in branch 10 compared to 9; right: genes upregulated in branch 9 compared to 10. **(h)** Dendogram showing URD inferred lineage tree of single cells captured at d1.5, sorted cells expressing eGFP (NKX2.5) or mCherry (TBX5) at d4.5, and d30 cells obtained by reaggregation of sorted cells on d4.5; differentiation with 0.5 μM RA for 4d (RA+). Arrows indicate the three main cell lineages recovered: cardiomyocytes (CMs), fibroblasts (FBs), and endothelial cells (ECs). Cells from each Seurat cluster are indicated by colour. **(i)** Quantification of flow cytometry analysis at d30 of cells expressing FN1 or cTnT within RA- differentiation or 0.5 μM 4d RA differentiation for 3 hPSC lines. Data are mean ± SEM; n = 3 differentiations per line. **(j)** Quantification of cells in myocytic clusters (TNNT2+) or non-myocytic clusters (FN1+) within d30 cells derived from CPC populations obtained during differentiation without (RA-) and with 0.5 μM RA for 4d (RA+). ECs: endothelial cells; FBs: fibroblasts; VCs: valvular cells; CPCs: cardiovascular progenitor cells, EndoPCs: endothelial/endocardial progenitor cells; CMs: cardiomyocytes; SMCs: smooth muscle cells; PCs: progenitors; MyoFBs: myofibroblasts; V-CMs: ventricular cardiomyocytes; OFT-CMs: outflow-tract cardiomyocytes; AVC-CM: atrioventricular canal cardiomyocytes; DEGs: differentially expressed genes.

Further commitment of the endocardial/endothelial cells towards a valvular-like cell fate was accompanied by downregulation of vascular endothelial markers such as *CDH5* (branch 23), which persisted in the endothelial like-cells of branch 9^32,33^ (Supplementary Fig. 7a). Subsequently, valvular-like cells divided into three branches (8, 10, and 11) based on the expression level of advanced EndoMT markers such as *POSTN* or *LUM*, and *BGN* which is expressed in the fibrosa layer of valves^33^ (Fig. 4d; Supplementary Fig. 7a).

By contrast, in the presence of RA, cardiovascular progenitors were already largely committed either to a non-myocytic (branch 19) or cardiomyocytic fate (branch 18) at day 4.5 (Fig. 4e), which is consistent with previous reports in the mouse suggesting limited potential of cardiac progenitors in the FHF^44^. The myocardial lineage trajectory was correlated with the sequential upregulation of *THBS4* and *WNT2* (Fig. 4f) resulting in several closely related ventricular-like CM branches (Supplementary Fig. 7b). In the non-myocytic branches, the cells rapidly differentiated either to an endothelial-like fate (branch 9; *CDH5*^+^, *CXCR4*^+^, *SOX17*^+^; Fig. 4g; Supplementary Fig. 7b; Supplementary Table 8) or to mesenchymal/fibroblast-like fate (branch 10, *THY1*^+^ *COL3A1*^+^ Fig. 4g; Supplementary Fig. 7b; Supplementary Table S8).

Notably, the fate decision trees of the different sorted populations (eGFP^+^, mCherry^+^, double mCherry^+^ eGFP^+^) arising from the CPC-RA+ were comparable, with similar marker expression throughout the trajectories (Fig. 4h; Supplementary Fig. 8a,b). This suggests that the different populations identified at day 4.5 in the presence of RA (eGFP^+^, mCherry^+^, double mCherry^+^ eGFP^+^) do not represent distinct progenitor pools but are more likely reflecting different stages of progression on the same differentiation trajectory. In addition, we found that the differentiation efficiency into the myocytic lineages, in both RA- and RA+ conditions, was similar between sorted (RA+:~87-96% cTnT^+^ CMs, ~4-12% FN1^+^ non-myocytic cell types; RA-: 84.7%; and 15.3%, respectively) and unsorted cells (RA+:~91-93% of cTnT^+^ CMs, ~5-10% of FN1^+^ non-myocytic cell types; RA-: ~83-92%; and ~13-17%, respectively), indicating that the negative (mCherry^-^ eGFP^-^) population present at day 4.5 does not seem to significantly impact the differentiation outcome (Fig. 4i,j). Taken together, these findings suggest that the differentiation protocol described in this manuscript can be utilized independently of the fluorescent marking of the hESC-line, providing a substantial advantage over previously described methods to isolate heart-field specific populations^45^.

### Human cardiac progenitors contribute to distinct cardiac structures in developing mouse hearts

To further investigate the lineage commitment and functionality of *in vitro* derived cardiovascular progenitors, we injected CPC-RA+ or CPC-RA- into the cardiac crescent region of developing mouse embryos, before initiation of visible contraction. These embryos were then subjected to whole-embryo *ex vivo* culture for 24 or 48 hours (Fig. 5a). The embryos’ heart developed normally, including beginning of contraction, formation of the linear heart tube, its looping and onset of chamber formation (Fig. 5b).

**Figure 5:**
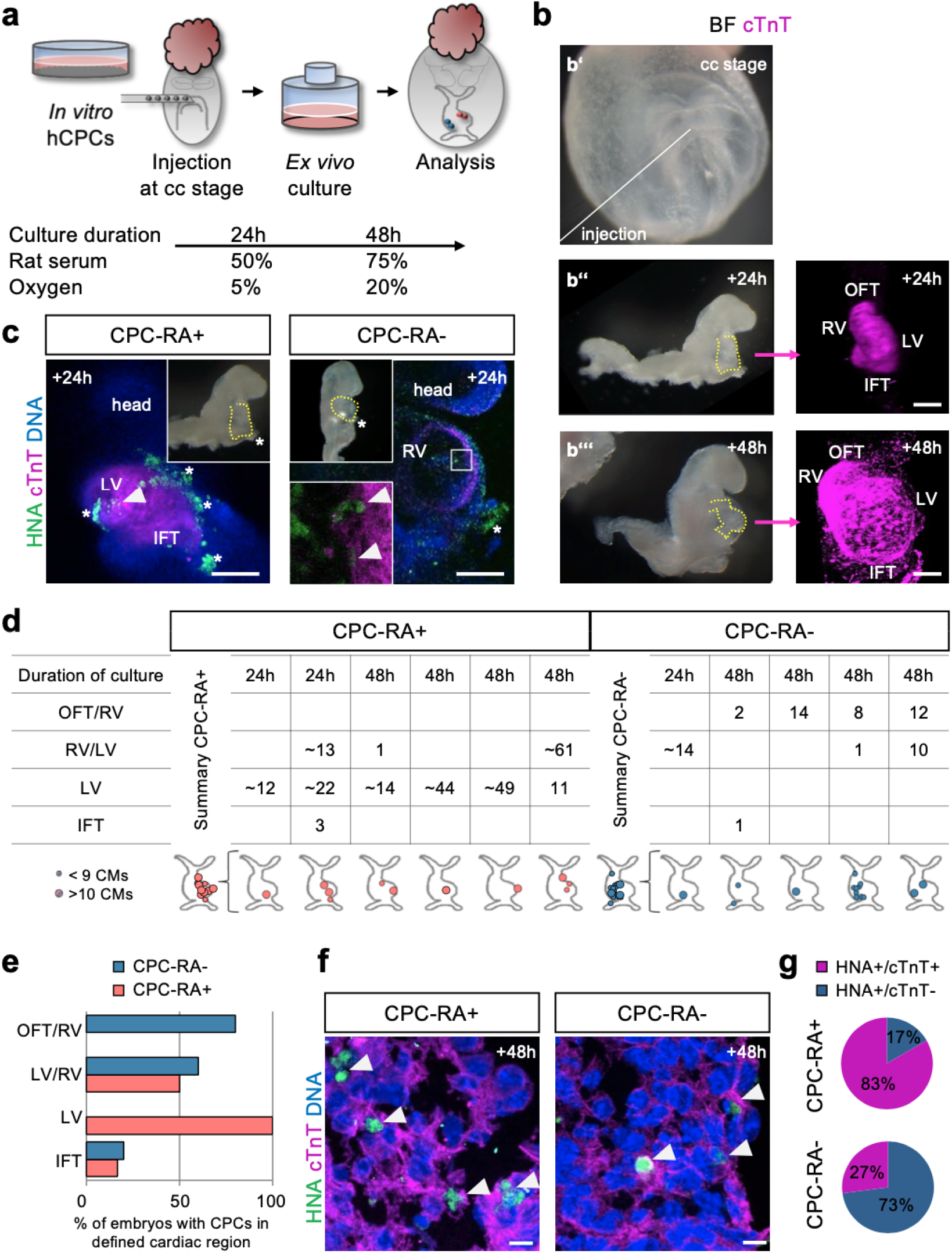
Human CPC-RA+ and CPC-RA- integrate within distinct regions of the developing mouse embryo heart. **(a)** Schematic representation of the protocol used to inject and culture murine embryos *ex vivo*. Rat serum and oxygen concentrations are increased after 24h of *ex utero* culture. **(b)** Representative brightfield images of cell injection into a cardiac crescent (cc) stage embryo (**b’**; the line indicates the site of injection) and after 24 and 48h of *ex vivo* culture (**b”** and **b”’**; dotted lines indicate the heart region). The right panels show a frontal 3D view of the heart of the embryos in b” and b”’ (marked by cardiac troponin, cTnT, in magenta), indicating elongation of the heart tube at 24h and looping at 48h. Scale bars = 100μm. **(c)** Representative confocal immunofluorescence images of the heart region (marked by cTnT) of embryos injected with human CPC-RA+ and CPC-RA- (marked by human nuclear antigen, HNA) after 24h of culture *ex vivo*. Scale bars = 100 μm. Top insets show an overview of the whole embryo. The bottom inset depicts a high magnification of the boxed region. Arrowheads indicate human cells within the murine heart. Stars indicate unspecific stain of the yolk sac. (**d)** Table and schematics summarizing the regional distribution and number of human cells derived from CPC-RA+ and CPC-RA- found integrated in the murine heart after 24 or 48h of *ex vivo* embryo culture. (**e)** Percentage of embryos with cells derived from CPC-RA+ or CPC-RA- in specific cardiac regions of the mouse heart after 24 or 48h of culture *ex vivo*. IFT: inflow tract; LV: left ventricle; OFT: outflow tract; RV: right ventricle. **(f)** High magnification images showing the integration of human CPC-derived cardiomyocytes (cTnT+) in the murine myocardium following injection of CPC-RA+ or CPC-RA-. Arrowheads indicate human cells marked by human nuclear antigen (HNA). Scale bars = 10μm. **(g)** Quantification of the percentage of human CMs (HNA+/cTnT+) and non-CMs (HNA+/cTnT-) found within the mouse heart following injection of CPC-RA+ or CPC-RA-.

The injected human cardiovascular progenitors from both conditions (CPC-RA+ and CPC-RA-) successfully integrated into the host myocardium (Fig. 5c-e; Supplementary Table 8). Whole-mount immunofluorescence analysis in cleared embryos showed that CPC-RA+ contributed mainly to the LV, with minor contributions to the inflow-tract (IFT; Fig. 5d,e). None of the injected embryos showed contributions to the OFT/RV region. Conversely, CPC-RA- contributed mainly to the OFT/RV, and we did not observe any of these cells in the LV (Fig. 5d,e). Immunofluorescence staining for cTnT indicated that cardiovascular progenitor cells from both conditions differentiated into CMs, with sarcomere organization of early myofibrils similar to the host CMs (Fig. 5f). Notably, the percentage of human non-myocytic cells (cTnT-negative) within the cardiac region was significantly higher after injection of CPC-RA- as compared to CPC-RA+ (Fig. 5g), most likely reflecting the broader lineage potential of the CPC-RA- population.

This data not only confirms our *in vitro* findings on the identity of CPC-RA+ and CPC-RA- at day 4.5 but also highlights that the cells at this stage are already committed to contributing to their respective cardiac structures.

### CPC-RA- and CPC-RA+ show disparate phenotypes in an *in vitro* model of hypoplastic left heart syndrome

To evaluate the utility of our fate-committed cardiac progenitors in studying human disease, we employed them as an *in vitro* model of HLHS, which is characterized by underdevelopment of the LV and the LVOT and is one of the most severe congenital heart defects^46^. We previously reported that HLHS patient-specific hiPSCs show defects in the differentiation of early cardiac progenitors resulting in abnormal cardiomyocyte subtype lineage specification and maturation^47^. However, the differentiation protocol used in that study had not allowed the investigation of heart field-specific differences.

Differentiating HLHS patient hiPSCs into FHF-like progenitors (CPC-RA+), we observed a significant downregulation of *NKX2.5* and *TNNT2* at d4.5 compared to healthy control hiPSCs (Supplementary Fig. 9a), which is in line with our previous findings on HLHS progenitors^47^. The reduction in *NKX2.5* levels was less pronounced when applying the differentiation conditions leading to aSHF-progenitors (*TNNT2* is not expressed at this stage in these conditions; Supplementary Fig. 9a). However, these cells showed a significant downregulation of *ISL1* and *BMP4* (Supplementary Fig. 9a).

At day 30, the HLHS patient-specific CMs arising from the CPC-RA+ displayed lower levels of ventricular cardiomyocyte transcripts (*MYH7*, *MYL2*) compared to control CMs, in particular the LV-specific marker *FHL2* (Supplementary Fig. 9b). At the same time, they exhibited higher expression levels of genes related to OFT-CMs (*RSPO3*) rendering their transcriptional profile similar to the patient-specific CMs arising from the CPC-RA- (Supplementary Fig. 9b). CMs from the aSHF-progenitors (CPC-RA-) displayed significant downregulation of markers associated with OFT-CMs, such as *RSPO3* (Supplementary Fig. 9b). This is likely a consequence of reduced *ISL1* and *BMP4* levels in the progenitors, since those genes are crucially required in the SHF for proper OFT formation in mice^37,48^.

The previously described maturation defect of HLHS patient-specific CMs^47^, reflected here by an increase in the *TNNI1/TNNI3* and *MYL7/MYL2* ratios compared to controls, was comparable between CMs derived from the CPC-RA+ and CPC-RA- (Supplementary Fig. 9c). This is consistent with observations *in vivo* showing that CMs from the LV as well as the RV of HLHS patient display maturation defects^47,49^.

Taken together, these findings suggest that HLHS patient-specific hiPSCs fail to acquire a FHF-like fate at the progenitor level, resulting in perturbed CMs without proper LVCM identity. Notably, they also show a defect in aSHF progenitor differentiation resulting in CMs with significant dysregulation of OFT-CM markers, providing a potential link to the LVOT defects observed in HLHS patients. This highlights how heart-field specific progenitors can increase the resolution of *in vitro* disease models and provide additional insights in human congenital heart disease.

## DISCUSSION

Due to the limited accessibility of early human embryonic tissue samples, our knowledge on the emergence of human cardiovascular progenitors and their contributions to the developing heart relies largely on findings from model organisms and *in vitro* model systems. In the current study, we describe a versatile *in vitro* differentiation platform that allows the generation and isolation of various cardiac progenitor pools from hPSCs by modulating RA-mediated signaling. RA signaling is one of the key signaling pathways in the developing embryo and is critically required for the organization of the trunk and the organogenesis of various tissues of all three germ layers, including the hindbrain, the eye, the spinal cord, the pancreas, and the heart^50,51^. It is first detected at late primitive streak stages in the paraxial mesoderm^52^ and plays an important role in defining the posterior boundary of heart fields^51,53^ as well as the proper specification of the pSHF that gives rise to the IFT^10^.

The latter has been exploited previously showing that addition of RA during *in vitro* differentiation of hPSCs towards CMs generates atrial CMs that are suitable for use in drug screening^12,22^. Notably, in our study we did not observe an upregulation of atrial CM markers after exposure to RA. Instead, the majority of cardiovascular progenitors found in our differentiation conditions had acquired a FHF-like fate rapidly giving rise to ventricular-like CMs. Differently to our approach, the aforementioned studies have either used an alternative cardiac differentiation protocol^22^, or applied RA later during differentiation^12^. In addition, the dosage of RA applied in our protocol is lower, and it has been reported previously that low doses of RA during hPSC differentiation improve differentiation efficiency but do not affect cardiac subtype specification^8,13^. Taken together this shows that timing and dosage of RA as well as the type of mesodermal progenitors exposed results in distinct differentiation outcomes highlighting the pleiotropic roles of RA during development^54^.

In addition to the “classical” first and second heart field progenitors, we identified a subset of cells that resembled progenitors of the recently discovered JCF, which contribute to the epicardium as well as myocardium in mice^5,6^. We uncovered that the surface protein ITGA8, which is expressed during development on epicardial cells in the mouse and the chicken^55,56^, can be used to isolate these cells using live cell sorting. Sorted JCF-like progenitors readily differentiated into epicardial as well as cardiomyocytic progeny, proposing the presence of a similar progenitor population during human cardiogenesis. Remarkably, this population was only present upon exposure to RA, suggesting that RA-mediated signaling might be critically involved in the generation of the JCF.

The toolbox presented in this study can be used to differentiate hPSCs into various mesodermal cardiac progenitor pools that contribute to heart formation, including aSHF, FHF, as well as the JCF, offering novel approaches for deciphering human cardiac development and for studying congenital heart disease. Combined with the current advances in self-organizing 3D cardiac tissue cultures^11,58–60^, it could provide the opportunity to generate even more complex heart structures and gain unprecedented insights into human cardiac development and disease.

## Author contributions

D.Z., A.G., A.B.M and A.M. conceived the study, interpreted the data, and wrote the manuscript. D.Z. designed and performed most of the experiments using hPSCs, including maintenance and differentiation of cells, cell sorting, single cell libraries preparation, and analyzed data. G.S. and A.G. performed bioinformatic analyses. J.K. performed mouse embryos injections, *ex-vivo* culture, and embryo analysis. D.O. and M.O. generated the ES03-TN line and provided conceptual advice. M.L. performed some molecular assays and immunostainings. T.D. supported analysis and data interpretation, and provided conceptual advice. S.C.H., M.N., A.T. performed patch clamp experiments. R.A.P. and P.G. provided conceptual advice. A.M., K-L.L. conceived and supervised the study and provided financial support. All authors read and approved the final manuscript.

## Acknowledgements

We would like to acknowledge Birgit Campbell, Christina Scherb, and Marco Crovella for their technical assistance, Gabi Lederer (Cytogenetic Department, TUM) for karyotyping, Dr. Rupert Öllinger (TUM, Germany) for sequencing, Dr. David Elliott for sharing the ES03 and ES03-NKX2.5^eGFP^ cell lines, Dr. Sasha Mendjan for advice and discussion, and Drs. Ed Stanley and Andrew Elefanty (MCRI, Australia) for advice in construct design and gene targeting.

## Data and materials availability

The scRNA-seq data will be made available at GEO database. All other data are available in the main text or the supplementary materials.

## Materials and methods

### hESC/hiPSC cell culture

Human iPSCs were generated using the CytoTune-iPS 2.9 Sendai Reprogramming Kit (Invitrogen; A16157) as previously described^47,61^. The following hiPSC lines were used in differentiation experiments: hPSCreg MRIi003-A (hiPSC/CTRL), MRIi020-A (HLHS). This study was approved by the Ethics Commission of the TUM Faculty of Medicine (# 447/17 S). Authorization to use the hESC line ES03 (hPSCreg ESIBIe003) generated by ES Cell International Pte Ltd in Singapore was granted by the Central Ethics Committee for Stem Cell Research of the Robert Koch Institute to AM (AZ 3.04.02/0131). Karyotype analysis was performed at the Institute of Human Genetics of the Technical University of Munich using standard methodology.

All hESC and hPSC lines were maintained in E8 medium containing 0.5% Penicillin/Streptomycin on Geltrex (A1413302, Gibco/Invitrogen) coated dishes under standard conditions (37°C, 5% CO_2_). Cells were non-enzymatically passaged every 4 days using 0.5 mM EDTA in PBS(-/-). To promote survival, Thiazovin was added at a concentration of 2 μM for 24 h after passaging.

### Generation of the reporter cell line by gene targeting

The method of gene targeting applied here was developed by the Stanley and Elefanty lab in Melbourne Australia (Costa et al. 2007; R. P. Davis et al. 2009). In brief, ES03 cells were grown on MEF feeders in KSR+bFGF and passaged enzymatically using trypsin-like enzyme (TrypLE, Invitrogen) for at least 3 passages before electroporation. 10 million cells were suspended in a total of 800 μl of ice-cold PBS containing 20 μg of linearised targeting vector in a 4mm gap cuvette. Electroporation was performed at 250 V and 500 μF. Cells were pipetted in pre-warmed medium and centrifuged at 259 g for 3 minutes. The cell pellet was resuspended in warm KSR containing 12 ng/ml bFGF and spread over nine 6-cm dishes containing antibiotic resistant MEF feeders. 5 days after plating, medium was changed to KSR+FGF containing G418. About 7 days after the start of selection, colonies were picked and split in two wells, a DNA well and a maintenance well. Cells in the DNA well were lysed for at least 3 hours at 55 °C (Lysis buffer: 100 mM TrisHCl, pH 8.0, 200 mM NaCl, 5 mM EDTA, pH 8.0, 0.2% (w/v) SDS powder (Sigma), 200 μg/ml proteinase K) and DNA was isolated using isopropanol precipitation. The DNA pellet was washed once in 70% Ethanol and after air-drying resuspended in 50 – 100 μl TE buffer. Long-range PCR over the 5’ homology arm was used to screen for correct integration.

The resistance cassette can potentially interfere with expression of the reporter gene (Pham et al. 1996; Scacheri et al. 2001). Therefore, removing the resistance cassette ensures that reporting of locus activity of the targeted gene is faithful and specific. The design we used for the targeting vectors includes rox sites flanking the puromycin resistance cassette. These rox sites can be recombined with Dre, the D6 site-specific DNA recombinase (Sauer & McDermott 2004), similar to the Cre/loxP system. To avoid issues with genomic integration of the expression plasmid, we chose to transduce the DRE protein directly using a fragment of the HIV TAT protein, which has been previously used as a cell-penetrating peptide (Frankel & Pabo 1988; Fawell et al. 1994). The TAT-DRE fusion protein was provided by Dr. Marko Hyvönen from the Department of Biochemistry (University of Cambridge). Cells grown on feeders were incubated with 2 μM TAT-DRE/CRE in serum free medium for 6 hours. After this treatment, the cells were placed back in KSR medium supplemented with bFGF. Removal of the resistance cassette was evaluated using primers flanking the resistance cassette. At the end of the process, cells were clonally derived by sorting single cells into 96 well plates in order to ensure a homogeneous population of targeted cells (and excision of the resistance cassette). Correct clones were identified by long range PCR on genomic DNA for targeting and the absence of the resistance cassette.

### Cardiac differentiation

Cardiac differentiation protocol was adapted from Mendjan et al., 2014, Hofbauer et al., 2021. Briefly, cells were plated on Geltrex-coated 24-well plates at a density of 200,000 cells/well in E8 medium containing 2 μM Thiazovin (day −1). The next day, medium was replaced with CMD-BSA medium – consisting of 1:1 DMEM/F-12 with Glutamax and IMDM containing 0.1 g/ml BSA, 30 mg/ml transferrin, 1% chemically defined lipid concentrate (11905031, Gibco), ~0.46mM (0.004%) of thioglycerol (T1753, Sigma) – supplemented with 10 ng/mL BMP4 (314-BP, R&D), 1.5 μM CHIR99021 (4423, R&D), 50 ng/mL Activin A (SRP3003, Sigma), 30 ng/mL bFGF (233-FB, R&D) and 5 μM LY 294002 hydrochloride (1130, R&D). After 40 h, the medium was replaced with CDM-Meso medium consisting of CDM-BSA supplemented with 10 ng/mL BMP4, 8 ng/mL bFGF, 10 μg/ml insulin (11376497001, Roche/Sigma), 5 μM IWP2 (72122, Stem Cells). Additionally, the medium was supplemented or not with RA (R2625, Sigma) in varying concentrations between 0.5 and 1 μM as indicated. CDM-Meso was changed the next 4 days in 24h intervals. Depending on the treatment, RA was added to media for 2 or 4 days. On day 5 and 6, the medium was replaced with CDM-BSA supplemented with 10 ng/mL BMP4, 8 ng/mL bFGF, and 10 μg/ml insulin. From day 7 on, the medium was replaced every second day with CDM-Maintenance medium consisting of CDM-BSA supplemented with 10 μg/ml insulin.

### FACS and flow cytometry analysis of live cells

For flow cytometry based sorting and analysis of live cells, cells between d0-d8 of differentiation were washed two times with PBS(-/-), dissociated with Accutase (A11105013, Gibco/Invitrogen 5min, 37°C), centrifuged at 1200 rpm for 5min, resuspended in 2% FCS in PBS, filtered through a 40 μm filter and subjected to sorting procedure on a FACS Aria III Cell Sorter (BD) and subjected to live flow cytometry analysis procedure on a Cytoflex S (Beckman-Coulter). DAPI (D3571, ThermoFisher Scientific) staining was used to discriminate dead and live cells (final concentration 0.01 ng/μl). Cells were sorted based on mCherry and eGFP expression and collected into 50-100% FCS in PBS, unless otherwise indicated. Analysis was performed using Kaluza software (Beckman-Coulter).

### 3D culture of sorted cells

After sorting, mCherry+, eGFP+, double eGFP+/mCherry+ or ITGA8+ cells were centrifuged, resuspended in CDM-BSA, counted, centrifuged again, and resuspended in CDM-Meso containing 0.5% Penicillin/Streptomycin, 10 μM of Rock Inhibitor Y-27632 and supplemented or not with RA in varying concentrations as indicated at a concentration of 100,000 cells per 200 μl per well of a U-shaped 96-well plate previously coated with 5% Poly(2-hydroxyethyl methacrylate. Plates were centrifuged for 2 min at 1,200 rpm and transferred to an incubator. From the next day on, the cardiac differentiation protocol was applied as described above. At d7 aggregates were transferred into the wells of a 48-well plate previously coated with 5% Poly(2-hydroxyethyl methacrylate) and put on a shaker. From this point on CDM-Maintenance was supplemented with 50 ng/μl VEGF (293-VE, R&D).

### Epicardial differentiation

For epicardial differentiation, d4.5 FACS sorted ITGA8+ cells were re-plated on 0.1% gelatin coated wells of 12-well chamber slides in density of 20 000 cells per 1 cm^2^ and subjected to modified protocol Bao et al.^62^. Briefly, cells were re-plated in LaSR medium consisting of Advanced DMEM/F12 containing Glutamax, Ascorbic Acid (0.1 mg/ml; A5960, Sigma) and 0.5% Penicillin/Streptomycin (LaSR) with addition of 1% FCS and 10 μM of Rock Inhibitor Y-27632. On d6 and d7 the medium was replaced with LaSR supplemented with 3 μM CHIR99021. From d8 onwards LaSR medium was replaced daily till d12.

### Single-cell dissociation

Cells were dissociated using Accutase up to day 8. From day 8 on cells or aggregates were subjected to papain-based dissociation. Briefly, cells/aggregates were washed 2 times with 2 mM EDTA. Dissociation was carried out using a papain solution prepared as described by Fischer et al.^63^. Cells/aggregates were incubated with papain solution for 20-40 min at 37°C (the optimal dissociation time was previously determined for each cell line/time point to obtain a single cell suspension without compromising cell quality and survival). Aggregates were incubated on a shaker at 37°C. After that time, a solution containing 1 mg/ml trypsin inhibitor (T9253, Sigma) was added. Cells/aggregates were dissociated by pipetting, transferred to a tube containing PBS(-/-), centrifuged at 1,200 rpm for 5 min and resuspended in appropriate solutions depending on the downstream analysis.

### RNA isolation, reverse transcription PCR (RT-PCR), and quantitative real-time PCR (qPCR)

For RNA isolation cells/aggregates were dissociated as described above. For time course analysis d0-d8 RNA collection was performed 2 hours after media change. In case of aggregates, 3 - 5 aggregates were collected from each differentiation for RNA collection. Cell pellets were lysed and RNA was isolated using the Absolutely RNA Microprep Kit (400805, Agilent Technologies), Absolutely RNA Nanoprep Kit (400753, Agilent Technologies) or Rneasy Microkit (74004, Qiagen), depending on the cell number. For RNA isolation of ITGA8+ sorted cells, after sorting and centrifugation in PBS, cell pellets of 20,000 cells were lysed. 0.3-0.5 μg of RNA was used to synthesize cDNA with the High Capacity cDNA Reverse Transcription kit (4368813, Applied Biosystems). 40 ng was used to synthesize cDNA from ITGA8+ cells with the SuperScript IV Vilo Master Mix (11756050, Invitrogen). Gene expression was quantified by qPCR using 1 μl cDNA, the Power SYBR Green PCR Master Mix (4367659, Applied Biosystems), the primers listed in Supplementary Table 11 and a 7500 Real Time PCR System (Applied Biosystems). Gene expression levels were quantified relative to *GAPDH* expression using the ΔCt method, unless otherwise indicated.

### Single-cell RNA sequencing (scRNAseq)

For scRNASeq at d1.5, cells were washed two times with PBS, dissociated with Accutase (3-5 min, 37°C), centrifuged at 1,200 rpm for 5 min, resuspended in 0.04% BSA in PBS, filtered through a 40 μm filter and counted. For scRNASeq at d4.5 of differentiation, cells were subjected to sorting as described above. Cells were sorted based on mCherry and eGFP expression and collected into 10% FCS in PBS. Then cells were centrifuged, washed in 0.04% BSA in PBS, centrifuged again, filtered through a 40 μm filter and resuspended in 0.04% BSA in PBS for counting. For scRNASeq at d30, aggregates were dissociated with papain as described above, filtered through a 40 μm filter, centrifuged at 1,000 rpm for 3 min and resuspended in 0.04% BSA in PBS for counting. After counting, 10,000 cells for each sample were processed using the Chromium Single Cell 3’ Library & Gel Bead Kit v3.1 (1000075, 10x Genomics), Chromium Single Cell B Chip Kit (1000073, 10x Genomics), and Chromium i7 Multiplex Kit (220103, 10x Genomics) to generate Gel Bead-In-EMulsions (GEMs) and single cell sequencing libraries. Libraries were pooled and sequenced using the NextSeq 500/500 (Ilumina; High Output v2 kit 75 cycles v2.5 flow cell) with 28 cycles in read1 for the 10x barcodes and UMIs in and 8 cycles i7 index read and 58 cycles for cDNA in read2 with a read depth of at least 20,000 pair reads per cell.

The Cell Ranger pipeline (v6.1.1) was used to perform sample demultiplexing, barcode processing and generate the single-cell gene counting matrix. Briefly, samples were demultiplexed to produce a pair of FASTQ files for each sample. Reads containing sequence information were aligned using the reference provided with Cell Ranger (v6.1.1) based on the GRCh37 reference genome and ENSEMBL gene annotation. PCR duplicates were removed by matching the same UMI, 10x barcode and gene were collapsed to a single UMI count in the gene-barcode UMI count matrix. All the samples were aggregated using Cell Ranger with no normalization and treated as a single dataset. The R statistical programming language (v3.5.1) was used for further analysis. Count data matrix was read into R and used to construct Seurat object (v4.0.1)^64^. The Seurat package was used to produce a diagnostic quality control plots and select thresholds for further filtering. Filtering method was used to detect outliers and high numbers of mitochondrial transcripts. These pre-processed data were then analysed to identify variable genes, which were used to perform principal component analysis (PCA). Statistically significant PCs were selected by PC elbow plots and used for UMAP analysis. Clustering parameter resolution was set to 1 for the function FindClusters() in Seurat. For sub-clustering analysis we used clustree package (v0.4.3).

All DEGs were obtained using Wilcoxon rank sum test using as threshold p-value ≤0.05. We used adjusted p-value based on Bonferroni correction using all features in the dataset. For the cell type-specific analysis, single cells of each cell type were identified using FindConservedMarkers function as described within Seurat pipeline. For all the gene signatures analysed we used a function implemented in *yaGST* R package (https://rdrr.io/github/miccec/yaGST/)^65^. To infer the developmental lineage relationships between cells in our study, we used the R package URD (version 1.1.1)^66^, which requires the user to declare the root samples of the cell lineage tree. URD algorithm traces routes through a cell-cell nearest neighbour graph producing a tree-graph that summarizes the lineage. We selected as root sample d1.5 for both generated URD cell lineage trees and d30 samples as the latest stage. Marker genes in each branch were defined using *markersAUCPR* function (adjusted p-val ≤0.05). Clusters enriched in cell cell-cycle related genes were removed from analysis.

### Flow cytometry

#### Intracellularly/intranuclearly stained cells

For intracellular/intranuclear staining for flow cytometry cells were dissociated, counted and distributed in equal numbers per sample in 15 ml tubes (typically 2×10^6^ cells per sample per tube). Cells were fixed with 4% PFA for 7 min at room temperature (RT; 500 μl/10^6^ cells), then centrifuged 3 min at RT at 400 g. Next cells were washed 3 times with PBS (shaked for 5 min and centrifuged 5 min at 400 g between each wash). Then cells were stored in 2% FCS in PBS+/+ at 4°C or incubated right away with blocking/permeabilization buffer containing 10% FCS, 0.1% Triton-X-100, 0.1% saponin in PBS(+/+) (for intranuclear staining) or 10% FCS, 0.1% saponin in PBS+/+ (for intracellular/membrane staining) for 1h at RT on a shaker (1 ml / 10^6^ cells). Then cells were centrifuged and incubated with the primary antibody (Supplementary Table 9) diluted in 1% FCS, 0.1% saponin with or without 0.1% Triton-X-100 in PBS+/+ (500 μl/10^6^ cells), overnight at 4°C on a shaker. Then cells were washed three times with 0.1% saponin in PBS+/+ with or without 0.1% Triton-X-100 for a total of 45 min on a shaker (cells were centrifuged between washes). Then the secondary antibody (Supplementary Table 10) was added diluted in 1% FCS, 0.1% saponin in PBS+/+ with or without 0.1% Triton-X-100, (500 μl / 10^6^cells) and cells were incubated for 1 hour at RT, protected from light, on a shaker. After that, cells were washed three times with 0.1% saponin in PBS+/+ with or without 0.1% Triton-X-100 for a total of 45 min on a shaker. Next, cells were resuspended in 2% FCS in PBS+/+ (100 μl / 10^6^ cells), passed through a 40 μm strainer and subjected to analysis on a CytoFlex S or Gallios (Beckman Coulter). Analysis was performed using Kaluza software (Beckman Coulter). No-primary antibody, nosecondary antibody, IgG antibody controls were performed.

#### Surface protein-stained cells

For flow cytometry analysis of surface protein-stained cells, cells were dissociated, counted and distributed in equal numbers per sample in 15 ml tubes (typically 5×10^6^ cells per sample per tube). Samples were incubated with ITGA8 or IgG-APC antibodies (Supplementary Table 9) diluted in the FACS buffer containing 2% FCS in PBS (10 μl antibody/100 μl buffer/10^6^ cells) for 30 min on ice. Then cells were washed three times with FACS buffer and resuspended in FACS buffer for sorting. Before sorting DAPI was added to samples at a final concentration of 0.01 ng/μl to discriminate dead and live cells. Cells were sorted into tubes containing 1 ml of 50% FCS in PBS, washed with PBS, centrifuged and resuspended in lysis buffer for RNA extraction or cell culture media for further differentiation.

### Immunofluorescence staining of cells

#### Cardiac cells cultured in monolayer

At d5 of cardiac differentiation, cells cultured in 12-well chamber slides (81201, Ibidi) were fixed with 4% PFA for 10 min at RT, washed 3 times with PBS and permeabilized with 0.1% Triton-X-100 in PBS for 15 min. After blocking with 10% FCS in 0.1% Triton-X-100 in PBS for 60 min, samples were stained using primary antibodies (Supplementary Table 9) diluted in 0.1% Triton-X-100 PBS containing 1% FCS overnight at 4°C The samples were then washed 5 times for 5min with 0.1% Triton-X-100 PBS and incubated with secondary antibodies (Supplementary Table 10) diluted 1:500 in 0.1% Triton-X-100 PBS containing 1% FCS for 1 hour at RT. Nuclei were detected with 1 μg/ml Hoechst 33258.

#### Epicardial cells

Cells at d12 of epicardial differentiation were fixed with 4% PFA for 15 min at RT, washed 3 times with PBS, blocked for 1 hour in 0.1% Triton-X-100 PBS containing 3% BSA and stained using primary antibodies (Supplementary Table 9) diluted in 0.1% Triton-X-100 PBS containing 0.5% BSA overnight at 4°C. Samples were then washed 3 times for 5 min with 0.1% Triton-X-100 PBS and incubated with secondary antibodies (Supplementary Table 10) diluted in 0.1% Triton-X-100 PBS containing 0.5% BSA for 1 hour at RT. Specimens then were washed 3 times with 0.1% Triton-X-100 PBS. Nuclei were detected with 5 μg/ml Hoechst 33258 (5 min). Slides were washed once again with PBS, covered with mounting medium and a cover slip and stored at 4°C.

### Mouse-human embryo chimeras

#### Timed mating and embryo dissection

All animal experiments were performed in accordance with German animal protection laws and EU ethical guidelines (Directive 2010/63/EU). The day after overnight mating, mice were separated and checked for the presence of a vaginal plug (ED0.5). On the desired day of embryonic development, the status of gestation was evaluated and pregnant females were sacrificed. Embryos were dissected out of the uterus and placed in a dish with pre-warmed dissection medium (5% FCS, 1% Pen/Strep, 20mM HEPES in DMEM). Embryos were carefully removed from the decidua and Reichert’s membrane. Attention was paid to not remove the ectoplacental cone or destroy the yolk sack. Damaged embryos were not used further.

#### Injection and whole embryo culture ex utero

CPCs were injected at d5 of differentiation into the cardiac crescent region of dissected mouse embryos. Approximately 20-50 cells per embryo were introduced *via* a glass capillary of 20 μm inner diameter. Operated and stage-matched unoperated embryos were placed into media containing 50% rat serum (S2150, Biowest) and 50% DMEM for 24h. Medium was changed after 24 hours to media containing 75% rat serum and 25% DMEM. Embryos were cultured using a rotating incubator system (BTC Engineering) at 5% CO_2_ and 5-20% O_2_ depending on their developmental stage. At the end of the culture, embryos were removed from the incubator, dissected from the yolk sac and amnion, and evaluated in terms of heart beat and overall morphological development.

#### Whole-mount immunofluorescence staining and clearing of embryos

After evaluation, properly developed embryos were transferred to tubes and fixed in 4% PFA at RT for 1-2h depending on size. Next, embryos were washed in PBS, and incubated in blocking/permeabilization solution containing 10% FBS, 0.1 % Triton X-100 in PBS on a shaker at RT for 4h. Embryos were then incubated with primary antibodies (Supplementary Table 9) diluted in 1% FBS, 0.1% Triton X-100 in PBS on a shaker at 4□°C overnight; after which they were 3x washed in 0.1% Triton X-100 in PBS for 3h in total; and incubated with secondary antibodies (Supplementary Table 10) and Hoechst dye diluted in 1% FBS, 0.1% Triton X-100 in PBS on a shaker at RT for 2h. After another round of washing the embryos were cleared using a protocol adapted from Masselink et al.^67^ Briefly, embryos were sequentially transferred into dehydration solutions containing 30%, 50%, 70 % and 2x 100% 1-Propanol in PBS (pH adjusted to 9.0-9.5 using trimethylamine) and incubated on a shaker at 4°C. Incubation times were adjusted considering the embryos’ size (at least 4 hours per dehydration step). After complete dehydration, the embryos were stored in ethyl cinnamate.

#### Immunofluorescence staining of cryosections

Embryos were subjected to a sucrose gradient (5-20%) followed by embedding in a 1:1 mixture of Tissue-Tek O.C.T. and 20% sucrose, and frozen in a bath of 2-methylbutane chilled with liquid nitrogen. Samples were stored at −80□°C or cryo-sectioned into 8□μm slices transferred to polysine-coated slides. Sections were fixed in 4% PFA at RT for 10 minutes. After washing with PBS, samples were incubated in solution containing 10% FBS, 0.1 % Triton X-100 in PBS at RT for 1.5 h. Next, they were incubated with primary antibodies (Supplementary Table 9) diluted in 1% FBS, 0.1% Triton X-100 in PBS at 4 °C overnight. The next day, samples were washed 3x with 0.1% Triton X-100 in PBS for 15 min total and incubated with secondary antibodies (Supplementary Table 10) and 5 μg/ml Hoechst 33258 diluted in 1% FBS, 0.1% Triton X-100 in PBS at RT for 1 h. After another round of washing, sections were covered with mounting medium and a cover slip and stored at 4°C.

### Confocal microscopy and image analysis

Cleared embryos were transferred into an 8 Well Glass Bottom μ-Slide (80827, ibidi) and imaged using confocal laser scanning microscopy (Leica Microsystems, SP8) and analyzed using the Leica Application Suite X. To quantify the contribution of human cells to the heart of injected embryos, HNA+ cells were counted and the region of integration was noted.

### Patch clamp electrophysiological recordings and data analysis

Cardiomyocyte monolayers were washed twice with PBS and dissociated to single cells by adding 0.25% Trypsin-EDTA (ThermoFisher Scientific (Gibco); 25200072) and incubating at 37 °C for 2-3 minutes. If clumps of cells were still present, the cells were incubated for a further 2-3 minutes. The cells were then pipetted gently six times to ensure dissociation and FBS was added to neutralize the activity of trypsin. Dissociated cells were centrifuged at 800 rpm for 3 minutes and resuspended in RPMI-1640 media (ThermoFisher Scientific (Gibco); 11875093) supplemented with Penicillin/Streptomycin (ThermoFisher Scientific (Gibco); 15140122), ITS-G (ThermoFisher Scientific (Gibco); 41400045), chemically defined lipid concentrate (ThermoFisher Scientific (Gibco) #11905031) and 1-Thioglycerol (Sigma Aldrich; M6145) (RI medium) containing 20% FBS. Cells were then seeded onto glass coverslips precoated with laminin (Sigma Aldrich; L2020). 24 hours later, cells were inspected for attachment and the medium was changed to RI medium. Cells were patch clamped 3-7 days after seeding (medium was changed every 3 days). Whole-cell current-clamp recording was carried out at room temperature (22 ± 2 °C) using an Axopatch 200B amplifier (Molecular Devices, USA). Pipette (Intracellular) solution contained (mM): 110 K-D-gluconate, 20 KCl, 10 NaCl, 2 EGTA, 10 HEPES, 1 MgCl2, 0.3 GTP, and 2 MgATP (pH 7.4 with KOH). Extracellular solution (Tyrode’s) contained (mM): 135 NaCl, 5.4 KCl, 5 HEPES, 1 MgCl_2_, 0.33 NaH_2_PO_4_, 2 CaCl_2_ and 10 Glucose (pH 7.4 with NaOH). Pipette resistance, when filled with intracellular solution, was ~2-3.5 MΩ. Action potentials were Liquid Junction Potential (LJP) corrected (LJPc). The LJPc (13.4 mV) was calculated using the *Clampex* Junction Potential Calculator (Molecular Devices, USA). AP (Action Potential) properties were analysed using *Clampfit* software (Molecular Devices, USA). Cardiomyocytes based on their action potentials were classified into 4 groups: Ventricular-like cardiomyocytes (V-CMs), Atrial-like cardiomyocytes (A-CMs); Nodal-like cardiomyocytes (N-CMs) and Intermediate (ventricular)-like cardiomyocytes (I-CMs) based on APD90/50 ratio. I-CMs = APD90/APD50 ratio between 1.4 and 1.8. V-CMs = APD90/APD50 ratio between 1.0 and 1.4. A-CMs = APD90/APD50 ratio > 1.8; N-CMs: were classified based on the typical shape of action potentials and signs of spontaneous depolarization.

### Statistics

Statistical analysis was performed with GraphPad Prism version 5 and 8 (La Jolla California, USA). Bar graphs indicate the mean ± SEM with all data points displayed separately, unless otherwise indicated. Data from two experimental groups were compared either by unpaired Student’s *t*-test or by Mann-Whitney-Wilcoxon test depending on the assumed distribution. For more than two experimental groups one-way analysis of variance (ANOVA) was used first, followed by Student’s *t*-test or by Mann-Whitney-Wilcoxon for groupwise comparisons in case of a statistically significant result from the overall analysis. A p-value < 0.05 was considered statistically significant, unless otherwise indicated.

