## Supplementary Material for "Retinoic acid signaling modulation guides *in vitro* specification of human heart field-specific progenitor pools"

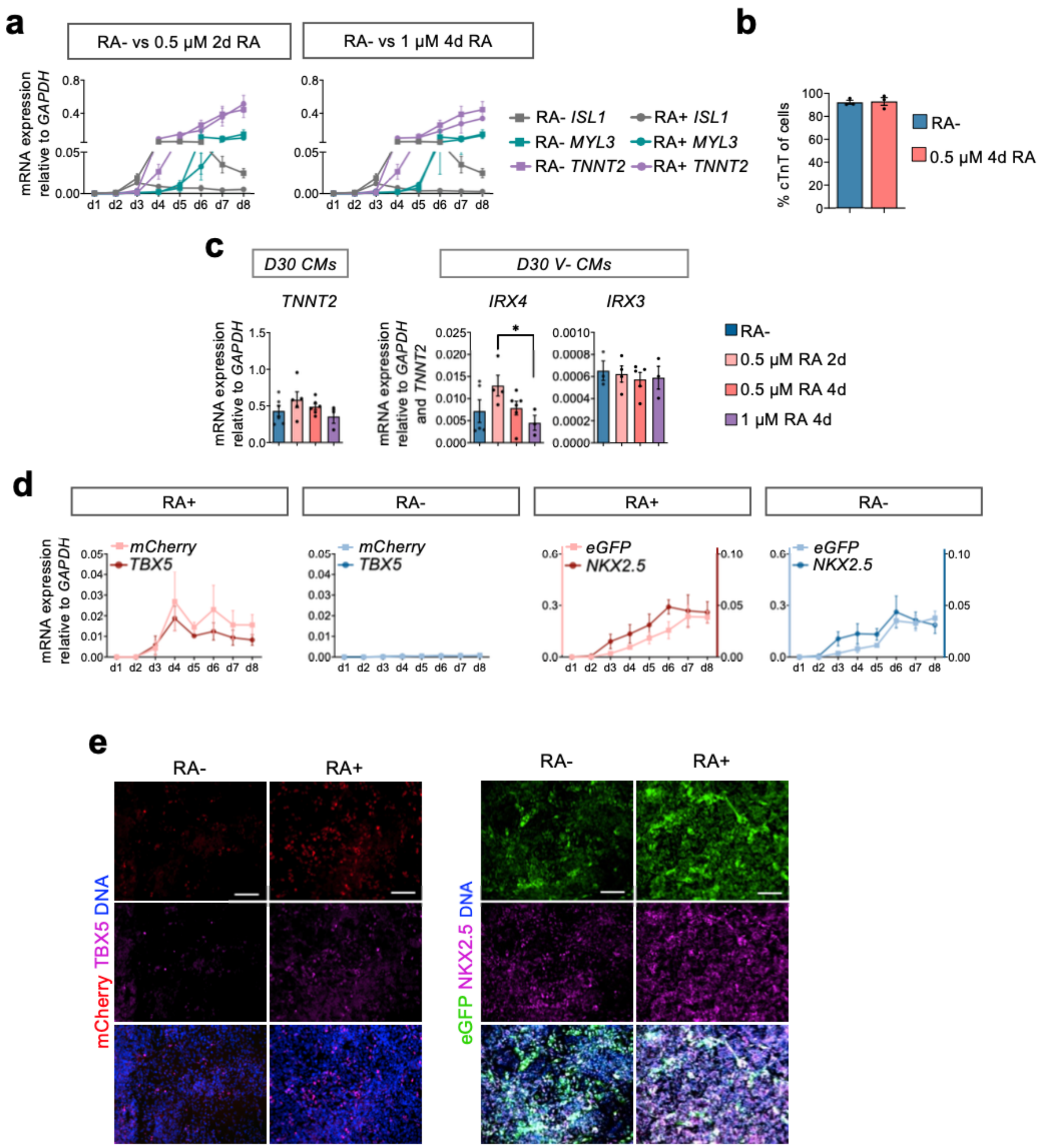


**Supplementary Figure 1: Generation of TBX5$^{mCherry}$ and NKX2-5$^{eGFP}$ hESC double reporter cell line to track the emergence of distinct pool of progenitors (related to Figure 1). (a)** Time course of mRNA expression of *ISL1* (CPCs) and *MYL3*, *TNNT2* (CMs) relative to *GAPDH* during differentiation without retinoic acid (RA-) compared to differentiation with different dosage and time of RA (RA+). Data are mean ± SEM. n ≥ 2 differentiations/time point. **(b)** Quantification of flow cytometry analysis of cells expressing cTnT at d30 of differentiation without retinoic acid (RA-) and with 0.5 µM RA for 4d. Data are mean ± SEM; n = 3 differentiations **(c)** mRNA expression of *TNNT2* and markers of ventricular cardiomyocytes (V-CMs; *IRX4*, *IRX3*) relative to *GAPDH* or *GAPDH* and *TNNT2*, respectively, at d30 of the indicated differentiation conditions. Data are mean ± SEM; n ≥ 3 differentiations *p<0.05

(unpaired two-tailed *t*-test). **(d)** Time course mRNA expression of *mCherry and TBX5* as well as *eGFP* and *NKX2.5* (relative to *GAPDH)* during differentiation without retinoic acid (RA-) and with 0.5 µM RA for 4d (RA+). Data are mean ± SEM; n = 3 differentiations **(e)** Representative immunofluorescence images of cells at d10 of differentiation without retinoic acid (RA-) and with 0.5 µM RA for 4d (RA+) stained with antibodies against mCherry (red) and TBX5 (magenta) (left) or eGFP (green) and NKX2.5 (magenta) (right). Nuclei were counterstained with DAPI (blue). Scale bars: 100 µm.

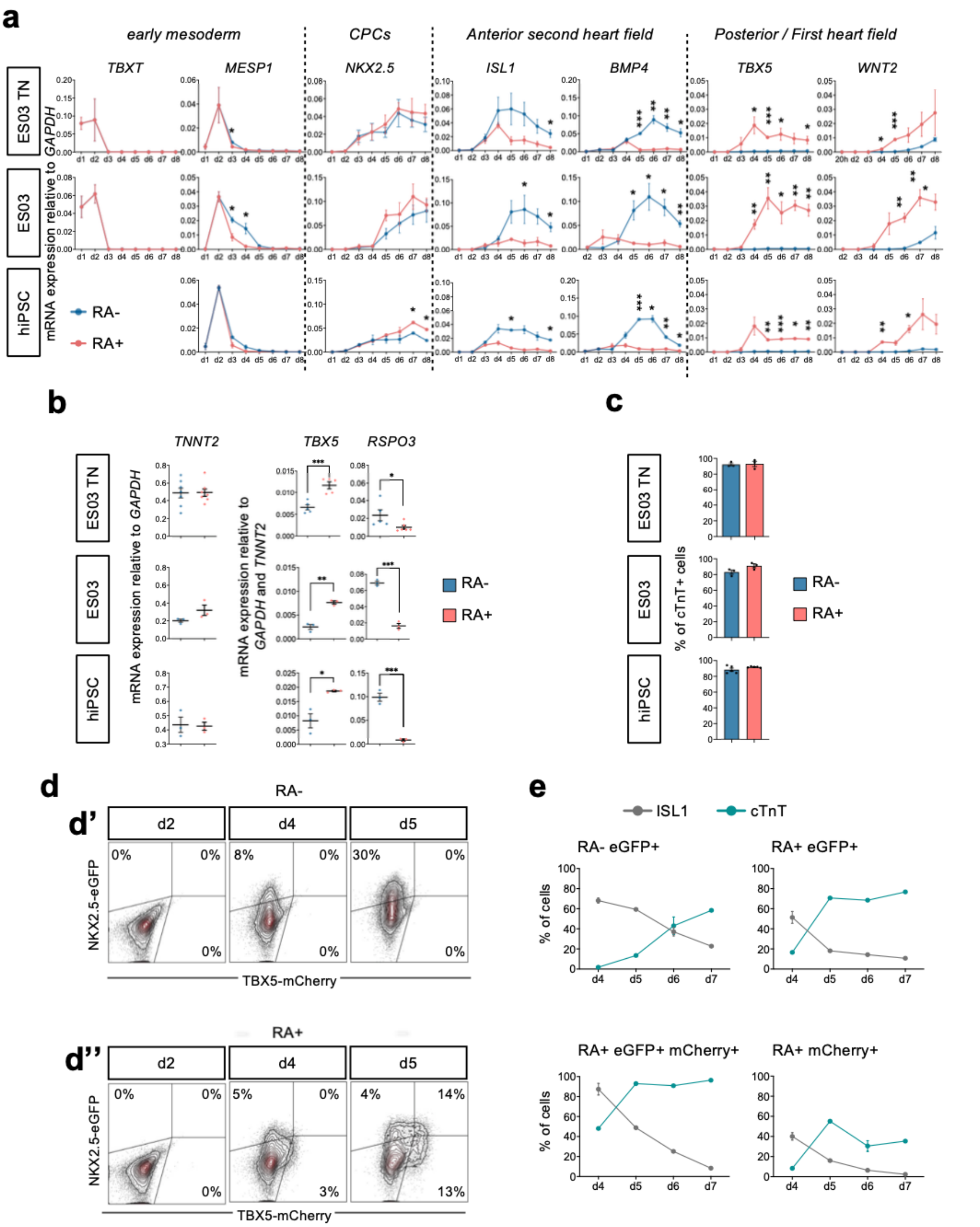


**Supplementary Figure 2: Cardiomyocyte differentiation is reproducible across human pluripotent stem cells lines (related to Figure 1). (a)** mRNA expression of key markers of early cardiovascular differentiation: *TBXT, MESP1* (early mesoderm), *NKX2.5, ISL1* (CPCs: cardiovascular progenitors); *BMP4* (aSHF: anterior second heart field), *TBX5* and *WNT2* (FHF: first heart field, posteriorization) relative to *GAPDH* of differentiation without retinoic acid (RA-) and with 0.5 µM RA for 4d (RA+) for 3 cell lines: ES03 TBX5-mCherry NKX2.5-eGFP (ES03 TN); parental line ES03 and independent hiPSC. Data are mean ± SEM; n ≥ 3 differentiations/time point for ES03 TN, n = 3 for ES03 and n = 2 for hiPSC. *$p<0.05$, **$p<0.005$,

***p<0.001 (unpaired two-tailed t-test). **(b)** mRNA expression of *TNNT2* (cardiomyocytes), *TBX5* (chamber cardiomyocytes), *RSPO3* (AVC, OFT cardiomyocytes) at d30 of differentiation without retinoic acid (RA-) and with 0.5 μM RA for 4d (RA+) for the 3 hPSC lines. Data are mean ± SEM; n ≥ 3 differentiations. mRNA expression relative to *GAPDH* for *TNNT2* or relative to *GAPDH* and *TNNT2* for other genes. *p<0.05, **p<0.005, ***p<0.001 (unpaired two-tailed t-test). **(c)** Quantification of flow cytometry analysis of cells expressing cTnT at d30 of differentiation without retinoic acid (RA-) and with 0.5 μM RA for 4d (RA+) for the 3 hPSC lines. Data are mean ± SEM; n = 3 differentiations/line. **(d)** Representative plots of live flow cytometry time course analysis of cells expressing mCherry (TBX5) and eGFP (NKX2.5) during RA- **(d')** and 0.5 μM 4d RA **(d'')** differentiation. **(e)** Quantification of flow cytometry time course analysis of cells expressing ISL1 and cTnT within mCherry$^+$/eGFP$^+$ (TBX5$^+$/NKX2.5$^+$); mCherry$^+$ (TBX5$^+$) and eGFP$^+$ (NKX2.5$^+$) populations from 0.5 μM 4d RA differentiation; and eGFP$^+$ (NKX2.5$^+$) population from RA- differentiation. Data are mean ± SEM; n ≥ 2 differentiations.

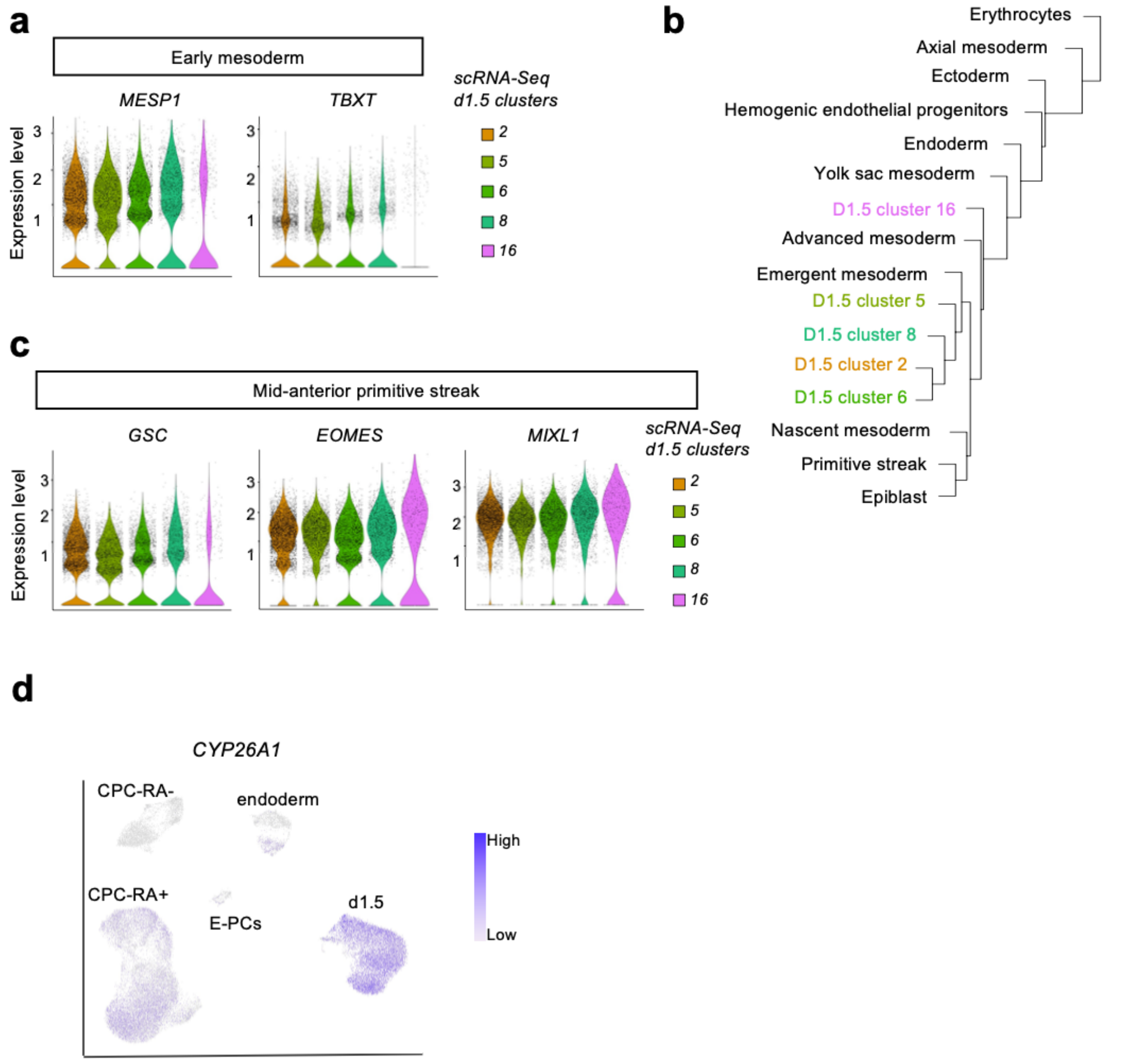


**Supplementary Figure 3: Cells at d1.5 correspond to mesodermal progenitors emerging from the mid-anterior primitive streak (related to Figure 2). (a)**. Violin plots showing the expression levels of early mesoderm genes (*MESP1*, *TBXT*) at d1.5 in the scRNA-seq clusters shown in Fig. 2b. **(b)** Dendrogram showing the integration of d1.5 scRNA-seq clusters with clusters from scRNA-seq analysis of *ex vivo* human gastrulation.[24] **(c)** Violin plots showing expression levels of genes defining the mid-anterior primitive streak stage (*GSC*, *EOMES*, *MIXL1*) at d1.5 in the scRNA-seq clusters shown in Fig. 2b. **(d)** Feature plot showing the expression of *CYP26A1* at d1.5 and d4.5 in the UMAP plot shown in Fig. 2b.

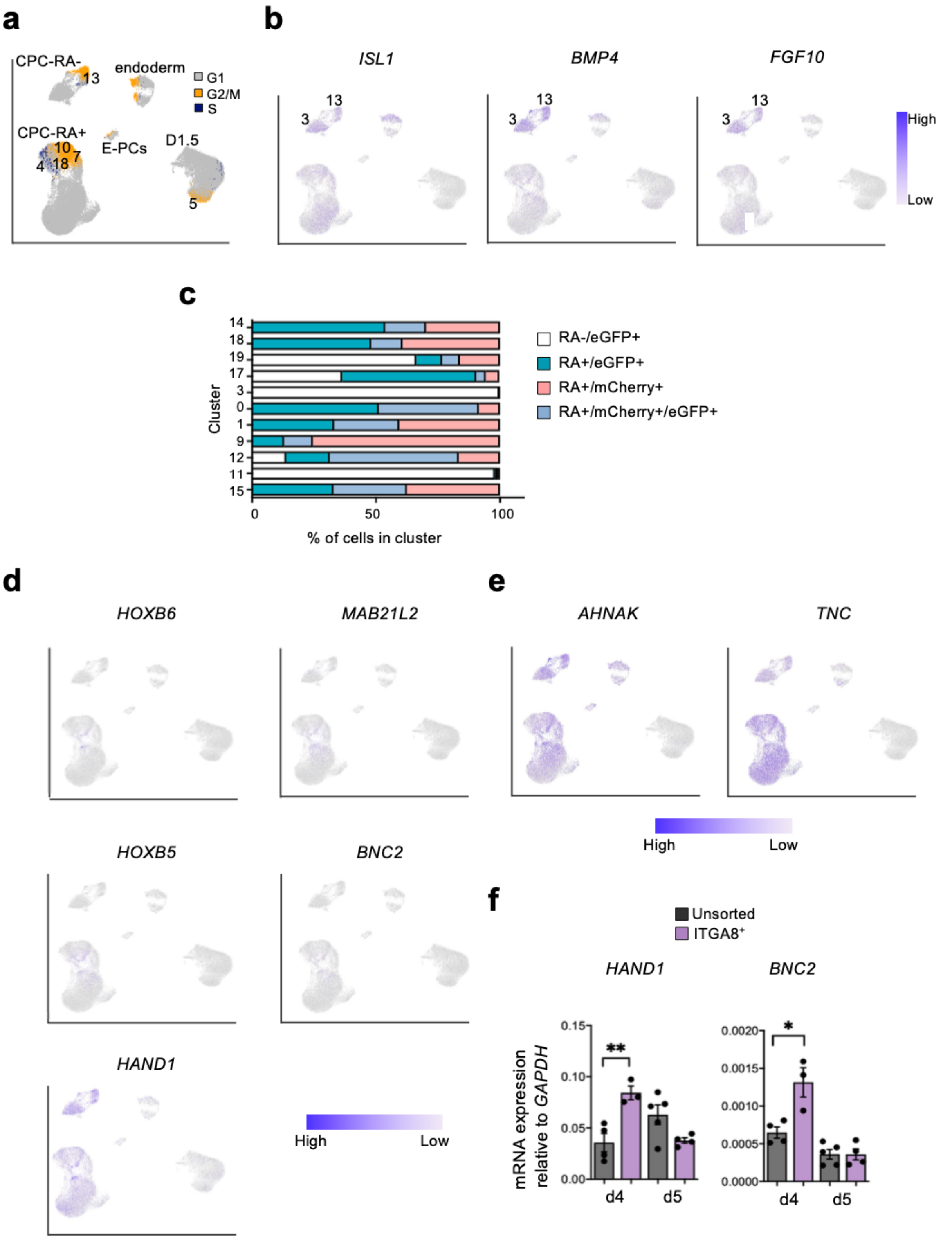


**Supplementary Figure 4: Characterization of human aSHF and JCF-like progenitors (related to Figure 2) (a)** Feature plot showing the expression of gene signatures specific of cell cycles phases – G2/M phase (yellow), S phase (blue), G1 (grey) – at d1.5 and d4.5 in the UMAP plot shown in Fig. 2b; main cell types are annotated. Clusters of proliferating cells are indicated. **(b)** Feature plots showing expression of key anterior second heart field (aSHF) markers (*ISL1, BMP4, FGF10*) in CPC-RA- clusters: 3 – non-proliferating; 13 – proliferating.

**(c)** Contribution of cells to the indicated clusters relative to total number of cells in the cluster. FACS sorted cells expressing mCherry (TBX5) and/or eGFP (NKX2.5) at d4.5 of RA+ (0.5 µM 4d) differentiation and eGFP (NKX2.5) at d4.5 of RA- differentiation. Clusters: 15, 11 - endoderm; 12 – early cardiovascular progenitors cells (CPCs); 9 – posterior second heart field (pSHF); 1, 0 – first heart field (FHF); 3 – anterior second heart field (aSHF); 17 – smooth muscle cell progenitor cells (SMC-PCs); 19 – endothelial/endocardial progenitor cells (E-PCs; 18, 14 – juxtacardiac field (JCF). **(d)** Feature plots showing expression levels of key JCF markers (*HOXB6, MAB21L2, HOXB5, BNC2, HAND1*) at d1.5 and d4.5 in the UMAP plot shown in Fig. 2b. **(e)** Feature plots showing the expression levels of two cell surface marker candidates (*AHNAK, TNC*) of JCF identified through comparative differential gene expression analysis between the mouse dataset of Tyser et al.[6] and our dataset. **(f)** mRNA expression of the JCF markers *HAND1* and *BNC2* relative to *GAPDH* in d4 and d5 cells FACS sorted for ITGA8 (APC) and unsorted cells during 0.5 µM 4d RA+ differentiation. Data are mean ± SEM; n ≥3 differentiations/time point; *p<0.05, **p<0.005, ***p<0.001 (unpaired two-tailed *t*-test).

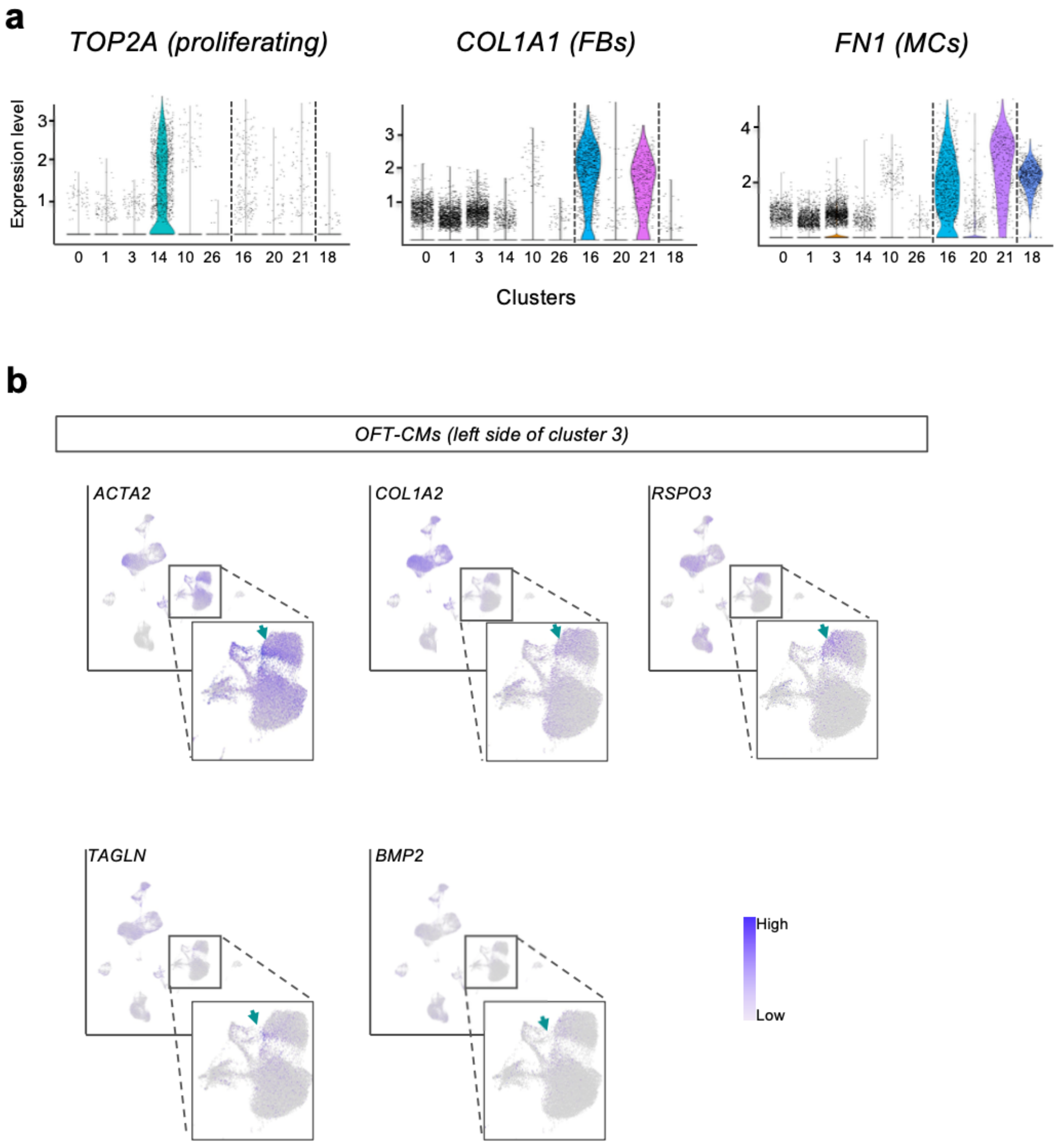


**Supplementary Figure 5: CPC-RA- and CPC-RA+ contribute to different cardiomyocyte subtypes clusters (related to Figure 3). (a)** Violin plots showing expression levels of the proliferation marker *TOP2A*, the fibroblast marker *COL1A1*, and the mesenchymal marker *FN1* in the scRNA-seq clusters shown in Fig. 3b. **(b)** Feature plots showing the expression levels of outflow tract cardiomyocytes (OFT-CMs) markers (*ACTA2, COL1A2, RSPO3, TAGLN, BMP2*) at d1.5, d4.5, and d30 in the UMAP plot shown in Fig. 3b. Insets highlight d30 cardiomyocyte clusters. Arrows indicate cells in the left side of cluster 3 identified as OFT-CMs.

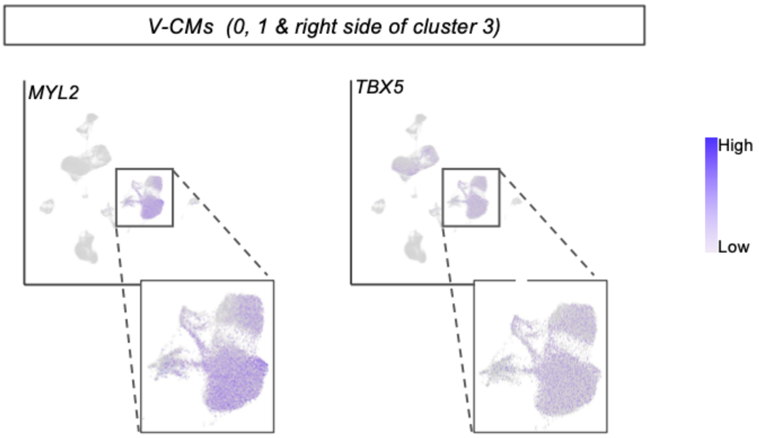


**Supplementary Figure 6: Both CPC-RA- and CPC-RA+ contribute to ventricular cardiomyocytes clusters (related to Figure 3).** Feature plots showing expression levels of ventricular cardiomyocytes (V-CMs) markers (*MYL2, TBX5*) at d1.5, d4.5 and d30 in the UMAP plot shown in Fig. 3b. Insets highlight d30 cardiomyocyte clusters.

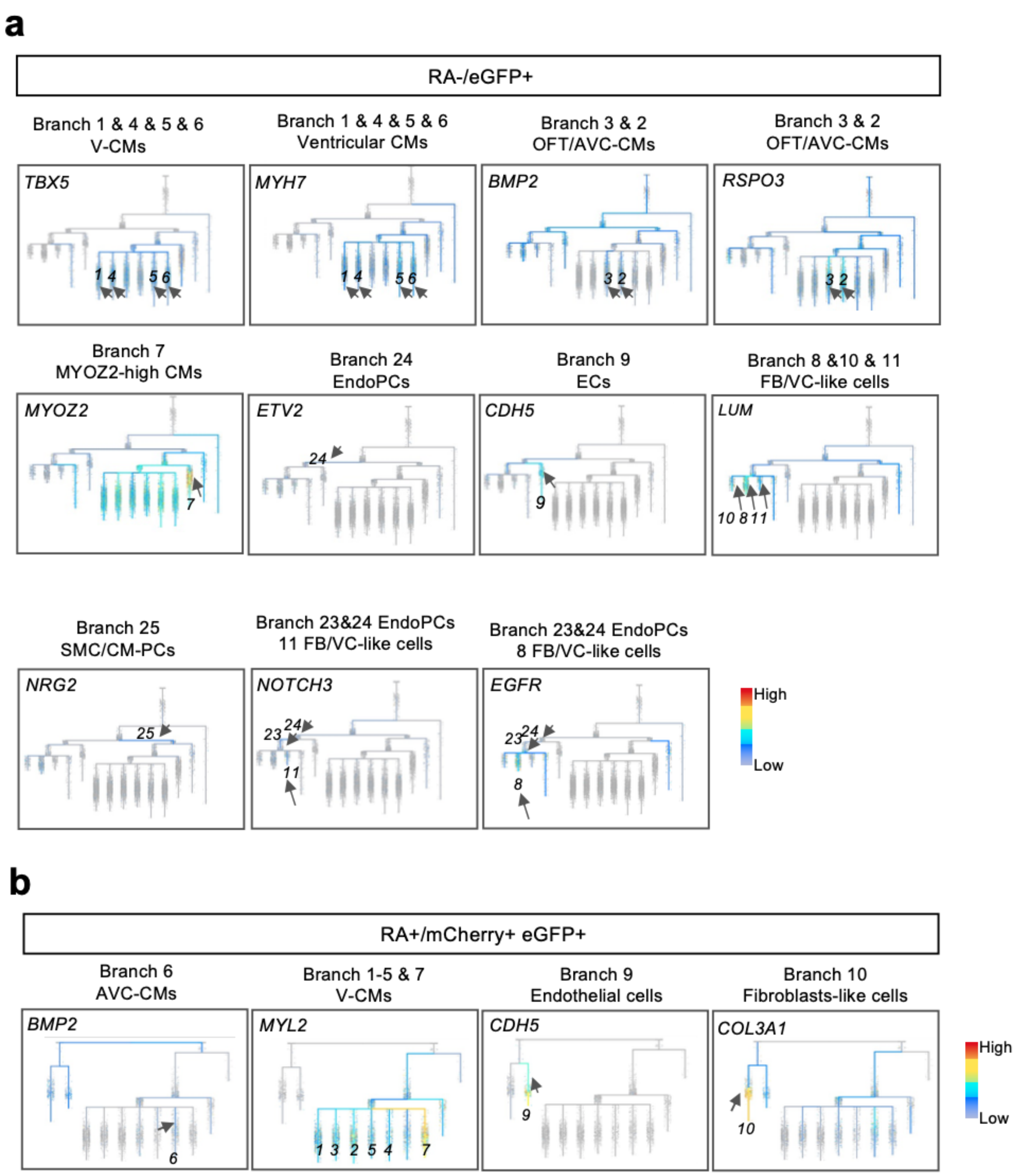


**Supplementary Figure 7: Differentiation trajectories uncover the specification stages of CPC-RA- and CPC-RA+ progenitors and their derivatives (related to Figure 4).** Marker genes differentially expressed among the lineages are plotted on the branches of the URD inferred lineage tree of single cells at d1.5, d4.5 (after FACS sorting), and d30 (after reaggregation of sorted cells). Cells were sorted for eGFP (NKX2.5) in RA- differentiation **(a)** and for eGFP (NKX2.5) and mCherry (TBX5) in RA+ differentiation**(b).** Arrows indicate branches with high expression of respective genes. CM: cardiomyocyte, OFT: outflow tract, AVC: atrioventricular canal, SMC: smooth muscle cell, FB: fibroblast, aSHF: anterior second heart field. VCs: valvular cells, MC: mesenchymal cells, V-CMs: ventricular cardiomyocytes, EndoPCs; endocardial/endothelial progenitor cells.

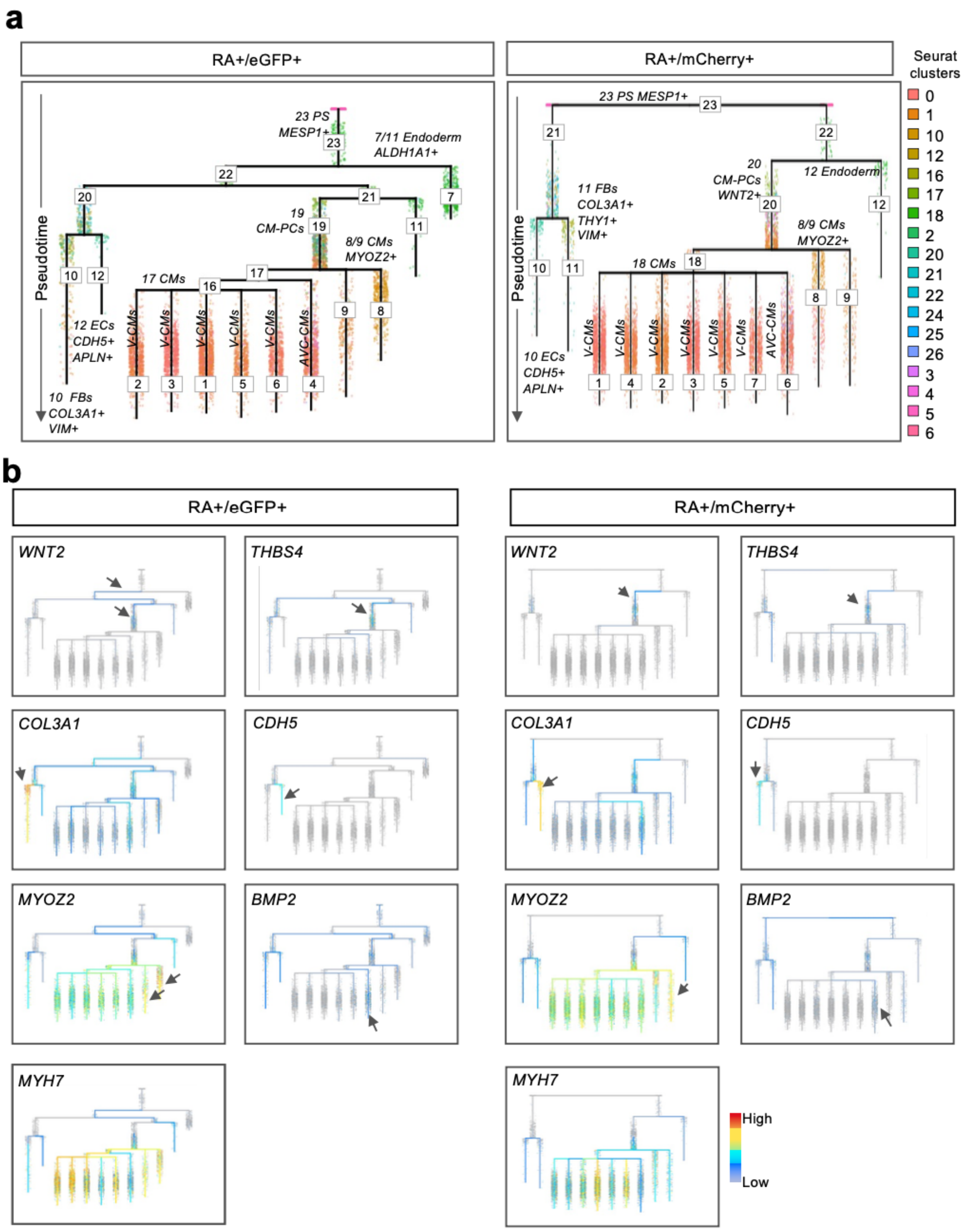


**Supplementary Figure 8: Differentiation trajectories uncover the specification stages of CPC-RA+ progenitors and their derivatives (related to Figure 4). (a)** Dendogram showing the URD inferred lineage tree of single cells at d1.5, d4.5 (after FACS sorting), and d30 (after reaggregation of sorted cells). Cells were sorted for eGFP (NKX2.5) in RA+ differentiation (left) or mCherry (TBX5) in RA+ differentiation (right). **(b)** Marker genes differentially expressed among lineages are plotted on the URD branches presented in (a). Arrows indicate branches with high expression of specific gene. CM: cardiomyocyte, OFT: outflow-tract, AVC:

atrioventricular canal, SMC: smooth muscle cell, FB: fibroblast, aSHF: anterior second heart field.

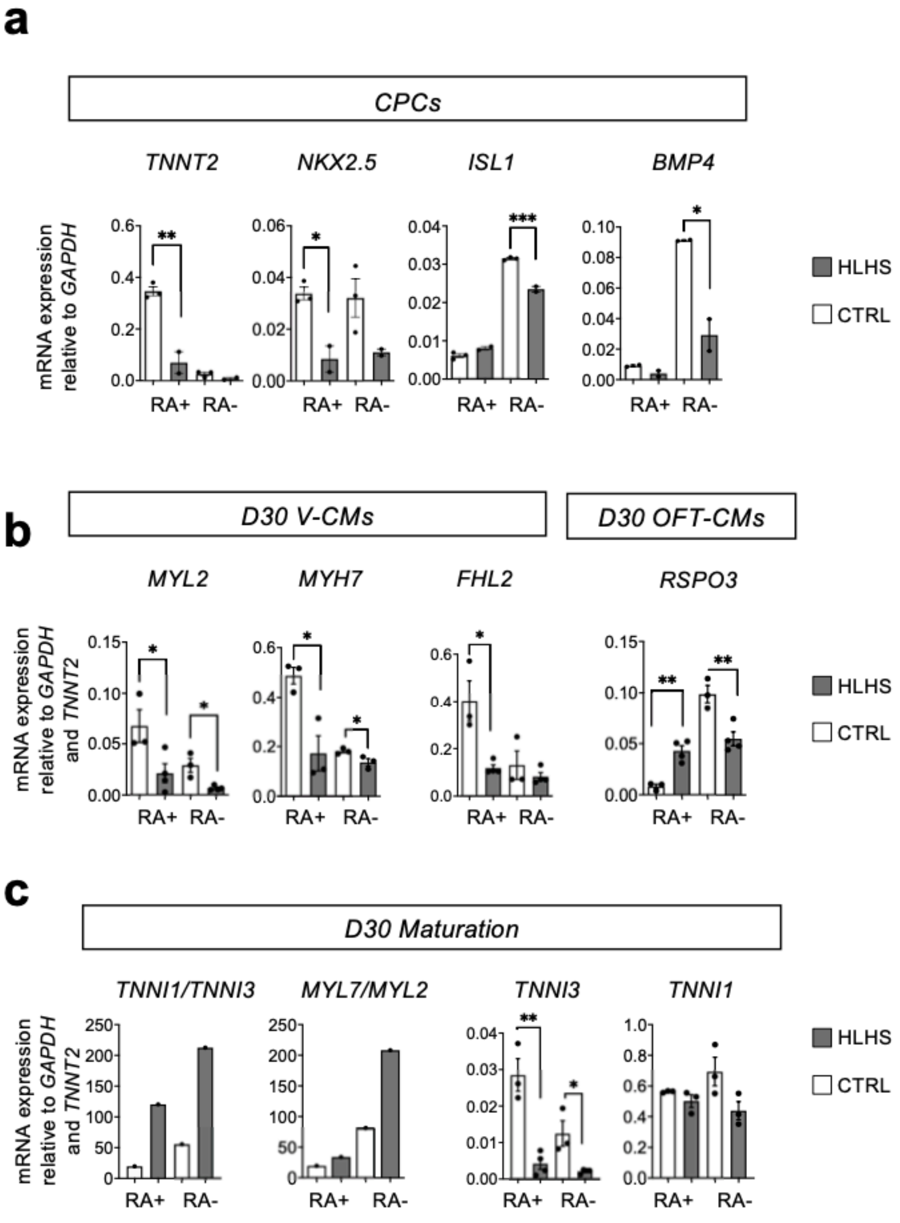


**Supplementary Figure 9: Pathological relevance of human heart field specification**. **(a)** mRNA expression of cardiovascular progenitor markers (*TNNT2, NKX2.5, ISL1, BMP4*) relative to *GAPDH* at d5 of RA- or RA+ (0.5 μM 4d RA) differentiation of HLHS patient-derived hiPSCs (HLHS) and healthy control hiPSCs (CTRL). Data are mean ± SEM; n ≥ 2 differentiations; *p<0.05, **p<0.005, ***p<0.001 (unpaired two-tailed *t*-test). **(b)** mRNA expression of markers of ventricular cardiomyocytes (V-CMs; *MYL2, MYH7, FHL2*), and outflow-tract cardiomyocytes (OFT-CMs; *RSPO3, BMP2, SOX4*) relative to *GAPDH* and *TNNT2* at d30 of RA- or RA+ (0.5 μM 4d) differentiation of HLHS patient-derived hiPSCs (HLHS) and healthy control hiPSCs (CTRL). Data are mean ± SEM; n ≥ 3 differentiations; *p<0.05, **p<0.005, ***p<0.001 (unpaired two-tailed *t*-test). **(c)** mRNA expression of maturation related markers (*TNNI1/TNNI3* and *MYL7/ML2* ratios, *TNNI3* and *TNNI1*) relative to *GAPDH* and *TNNT2* at d30 of RA- or RA+ (0.5 μM 4d) differentiation of HLHS patient-derived hiPSCs (HLHS) and healthy control hiPSCs (CTRL). Data are mean ± SEM; n ≥ 3 differentiations; *$p<0.05$, **$p<0.005$, ***$p<0.001$ (unpaired two-tailed *t*-test).

### Supplementary Tables

**Supplementary Table 1. (separate file)**

Parameters of action potential traces. Related to figure 1.

**Supplementary Table 2. (separate file)**

List of differentially expressed genes from the combined analysis of d1.5 and d4.5. Related to figure 2.

**Supplementary Table 3. (separate file)**

List of differentially expressed genes used to identify surface markers for the JCF. Related to figure 2.

**Supplementary Table 4. (separate file)**

List of differentially expressed genes from the combined analysis of d1.5, d4.5, and d30. Related to Figure 3.

**Supplementary Table 5. (separate file)**

List of differentially expressed genes from the subclustering of the non-myocytic cells from the combined analysis of d1.5, d4.5, and d30. Related to Figure 3.

**Supplementary Table 6. (separate file)**

List of differentially expressed genes from the URD analysis of CPC-RA-. Related to Figure 4.

**Supplementary Table 7. (separate file)**

List of differentially expressed genes from the URD analysis of CPC-RA+. Related to Figure 4.

**Supplementary Table 8. (separate file)**

Summary of mouse embryo experiments.

**Supplementary Table 9. (separate file)**

Primary antibodies used for immunofluorescence/FACS staining.

**Supplementary Table 10. (separate file)**

Secondary antibodies used for immunofluorescence/FACS staining.

**Supplementary Table 11. (separate file)**

Sequences of primers used for qPCR.
